# A Mechanistic Whole Brain Model to Capture Simultaneous EEG-fMRI Data

**DOI:** 10.1101/2025.07.23.666380

**Authors:** Anirban Bandyopadhyay, V. Srinivasa Chakravarthy, Dipanjan Roy

**Affiliations:** Computational Neuroscience Lab, Biotechnology, Indian Institute of Technology Madras, Play Field Ave, 600036, Chennai, India; School of Artificial Intelligence and Data Science (AIDE), Indian Institute of Technology Jodhpur, N.H. 62, Nagaur Road, 342030, Jodhpur, India

**Keywords:** Simultaneous EEG-fMRI, Hebbian Learning, Hopf Oscillator, Functional Connectivity, Whole Brain Model

## Abstract

This study introduces a novel oscillatory network model to simulate simultaneous EEG-fMRI data, addressing the reconstruction challenge that arises due to their contrasting spatiotemporal scales. Here, each brain region is modelled by two oscillator clusters - a cluster of low-frequency (LFO) and high-frequency Hopf oscillators (HFO) coupled with an innovative power-coupling rule, facilitating cross-frequency interactions. The model is trained in two stages: learning oscillators’ frequencies and phase relations using a biologically plausible complex-Hebbian rule in the first stage and, followed by a modified complex backpropagation for amplitude approximation, overcoming limitations of poor accuracy and computational complexity in existing models. This framework outperforms current methods in replicating empirical Functional Connectivity (FC), Functional Connectivity Dynamics (FCD), and modularity over disparate spatio-temporal scales. The correlation between the FC of fMRI and the FCs of various EEG frequency bands is reflected in the strengths of the LFO-HFO coupling. Furthermore, in-silico structural perturbation studies quantified the effect of pruning of the anatomical connectivity on spatiotemporal dynamics in terms of FC, FCD, modularity, and integration level (ISOR). The model’s ability to reconstruct simultaneous EEG-fMRI data showcases significant advancement in understanding the resting-state brain’s functionality from multimodal settings and deciphering neurological disorders in diverse spatiotemporal scales.

## Introduction

Recent advancements in neuro-imaging techniques allow neuro-scientists to systematically decipher various timescales associated with large-scale brain network dynamics. One of the most commonly employed methods for acquiring neuronal dynamics at a faster timescale is electroencephalography, or EEG for short. EEG uses electrodes over the scalp constrained by the skull geometry to acquire data that reveals the patterns of electrical activity of an ensemble of excitatory neurons located beneath the scalp. While EEG is highly temporally resolved (in the order of milliseconds) at revealing electrical activity and synchronization across the scalp’s surface, it is less effective at linking the observed activity to specific locations (poor spatial resolution) in the brain. Another widely used technique is functional magnetic resonance imaging or fMRI. The fMRI allows the detection of changes in the level of oxygen at different brain regions to measure how the activity of these regions changes over multiple seconds (ultraslow process). In contrast to EEG, fMRI is good at pinpointing the location of brain activity, but it is an indirect measure of brain activity as it depends on blood flow and several other factors. In terms of understanding how the brain works, EEG and fMRI thus provide different pieces of the puzzle. But there is no easy way to fit these pieces together.

Since the last two decades, there has been a surge of experimental studies using simultaneous EEG-fMRI, leveraging the strengths and deficits of each method. Combining these complementary methods has the potential to provide crucial insight into brain function that cannot be measured by one modality alone. Recent works along this line have shown promising results and gained deeper insights into understanding neurological disorders like-Epilepsy, various sleep stages, and a myriad of cognitive functions (Warbrick, 2022; Jorge et al., 2014; Balsdon et al., 2024).

Here we outline three related questions regarding understanding brain dynamics with simultaneous EEG-fMRI study. Firstly, a comprehensive understanding of the brain-dynamics is possible by measuring the functional activity of the brain by evaluating the linear or non-linear relationship between the parcellated brain regions’ neural activity (referred to as functional connectivity, FC) and the physical interconnections between them constrained by white fibre tracts (referred to as structural connectivity, SC). The manifestation of functional connectivity by structural connectivity, as supported by multiple studies, makes it evident that its aberration can cause several neurological disorders (Babaeeghazvini et al., 2021). To this end, a mathematical framework supported by empirical data can be an ideal candidate for deciphering the relationship between SC and the functional patterns recorded from individual or simultaneous EEG-fMRI. Most existing mathematical models have focused on either EEG or fMRI, relying on the conventional approach of optimizing parameters that govern the dynamics of the local and global neural activity to approximate the empirical data of EEG and fMRI (Schirner and Ritter, 2023; Wischnewski et al., 2022; Glomb et al., 2022). However, these parameter optimization processes are often time-consuming and computationally expensive, and the necessary parameters can vary across different empirical datasets. Moreover, such models have poor accuracy (in terms of matching simulated and empirical FC) in modelling EEG or fMRI (even both), which fails the purpose of building a whole brain model for mechanistic understanding.

Secondly, few recent works have built frameworks to capture simultaneous EEG-fMRI data; however, similar to the models built exclusively for either EEG or fMRI, these models also exhibit similar shortcomings: poor accuracy and rigorous parameter optimization. Moreover, such models mostly employ canonical haemodynamic response function (HRF) of a particular size and shape (e.g. fixed amplitude or response height, time to peak, full-width at half max or FWHM) to decode the hemodynamic activity from cortical neural activity and the popular Ballon-Windkessel model (Friston et al., 2000) to predict the BOLD signals without taking into account the variability of size and shape of HRF for different ROIs. The size and shape of HRF vary between different ROIs within a single individual (Devonshire et al., 2012; Handwerker et al., 2004; Rangaprakash et al., 2018); for example-the temporal lobe and orbitofrontal regions elicit higher HRF differences compared to other ROIs (Rangaprakash et al., 2018, 2023). Similar substantial variability has been observed between subjects and even between sexes, ages and eventually, such HRF variability or HRFv causes erroneous estimation of FC (West et al., 2019; Rangaprakash et al., 2023). Rangaprakash et al. found that the HRFv can alter the FC up to 50%, and the HRFv has an exponential relationship with the variability/ alteration of FC due to HRFv (Rangaprakash et al., 2018). Hence, adopting standardized canonical HRF can lead to misleading functional connectivity and confound the inference from the data and model. For instance, it can exacerbate the inaccuracy of the parameter optimization process because of the incorrect assumption (like-amplitude mismatch and temporal difference) of the HRF function pertaining to a particular ROI. On the other hand, a phenomenological model can suffice if it can produce sufficient mechanistic inference about the brain dynamics from the model without considering the detailed biological neurovascular coupling event.

Thirdly, the integrated EEG-fMRI models developed so far have not been intended to capture the relationship and overlap between the FC maps derived from both low-frequency oscillations of BOLD signals and the fast-frequency oscillations of EEG signals. Although many studies have aimed to understand physiological and cognitive processes using FC maps from EEG and fMRI, the neural basis of hemodynamic signals remains underexplored. The frequency-specific cross-modal relationship between EEG/MEG and fMRI has been explored in a few earlier empirical works and shows a statistically significant match between the FC map from fMRI and FC derived from EEG (Wirsich et al., 2020, 2021; Hipp and Siegel, 2015). Also, the frequency-specific (e.g. alpha, beta, delta, gamma, and theta bands) and positive correlation between the FCs from two modalities can reveal insights into information processing between neural activity and hemodynamic signals for the resting state condition (Wirsich et al., 2020). However, the underlying neurophysiological mechanisms of how neurons and blood vessels interact with each other are still an open question. In the context of neurological disorders like Epilepsy, where concurrent EEG-fMRI has been routinely used to deduce the epileptogenic network, simultaneously taken EEG-fMRI relationship reveals that the correlation between EEG-FC and fMRI-FC alters for temporal lobe epileptic patients, and it impacts the differently for right and left temporal lobe Epilepsy, elucidating that the temporal lobe epilepsy differently impacts the left and right parts of the brain, or in short, causing lateralization (Wirsich et al., 2024). Although empirical studies have shown the relationship between EEG-FC and fMRI-FC with structural connections, very few modelling studies have attempted earlier to integrate the three modalities. In the current modelling study, such an effort has been made to simultaneously capture the EEG and fMRI signals from all the ROIs, which are from similar ATLAS or spatially co-registered and provide mechanistic insights into the cross-modal, frequency-specific relationship.

To this end, different modelling and data driven inference approaches have emerged: EEG-driven fMRI prediction, fMRI-informed EEG estimation, comparison of concurrent EEG-fMRI (Warbrick, 2022; Lei, 2019). For fMRI-informed EEG estimation, spatio-temporal information obtained from fMRI is leveraged to guide the source reconstruction of EEG signals. In contrast, EEG-driven fMRI estimation has been used to predict the BOLD signal utilizing the canonical HRF (Hemodynamic Response Function), which acts as a convolution function and characterizes the temporal delay between EEG and BOLD signal. For comparison between EEG-fMRI, preferably, MRI-constrained source reconstructed EEG signal for each ROI is compared with fMRI data evaluated for each ROI, and it’s mostly the comparison between the functional connectivity map generated by each modality (Wirsich et al., 2021; Hipp and Siegel, 2015). However, the neural basis of the fMRI signal is still not fully understood from the empirical data itself. Functional activity generated from the EEG and fMRI do not fully match each other, rendering the opportunity to build a mathematical framework to simultaneously capture obtained EEG-fMRI data and infer the relationship between the signals with different temporal features.

In the extant literature, Large-scale brain network models are proposed to explain the underlying neural mechanisms behind the synchronization in networks of neurons, spontaneous oscillatory activity, excitation-inhibition balance, myelination and network topology (Chakraborty et al., 2025; Deco et al., 2017b, 2021; Vohryzek et al., 2022; Vattikonda et al., 2016; Pathak et al., 2022a); few attempts have been made in the realm of electrophysiological data (EEG/MEG) (Cabral et al., 2014; Pathak et al., 2022b; Glomb et al., 2020; Deco et al., 2017a). large-scale brain network models proposed earlier are systems of coupled neural mass models for simulating large-scale brain activity; coupling is often mediated by estimations of the strengths of anatomical connections based on diffusion-weighted MRI data (so-called structural connectivity (SC) or ‘connectomes’). Various models have attempted to elucidate the mechanisms underlying each modality, such as-model developed by Glomb et al.(2022) for EEG (Glomb et al., 2020); Abeysuriya et al.(2018), Cabral et al.,(2022), Hadida et al.(2018), Raj et al.(2020), Tewarie et al.(2019), Pathak et al. (2022) for MEG (Abeysuriya et al., 2018; Cabral et al., 2022; Hadida et al., 2018; Raj et al., 2022; Tewarie et al., 2019; Pathak et al., 2022b); Atasoy et al.(2016), Deco et al. (2017, 2021), Surampudi et al. (2018), Honey et al.(2007), Roberts et al.(2019), for fMRI (Atasoy et al., 2016; Deco et al., 2021, 2017b; Surampudi et al., 2018; Honey et al., 2009; Roberts et al., 2019). However, the characterization of features detected across modalities by large-scale brain network modeling approaches has only recently been initiated (Schirner et al., 2018; Rabuffo et al., 2021). In particular, Rabuffo et al. (2021) demonstrated how neuronal cascades can be a major determinant of spontaneous fluctuations in brain dynamics captured with simultaneous EEG and fMRI. Nonetheless, the relationship between large-scale oscillations and their organization across scales is yet to be thoroughly explored. Among the fewer attempts made for modelling multimodal imaging data, some proposed models exhibit several shortcomings: poor accuracy and rigorous optimization as stated earlier. (Schirner and Ritter, 2023; Wischnewski et al., 2022). Also, the models adopted so far cannot conjure the relationship and overlap between the low-frequency oscillations of BOLD signals and the fast-frequency oscillations of EEG signals. On the other hand, the model described in the current study deploys complex Hebbian learning to decipher the relationship between fast-spiking EEG and ultraslow fMRI signal. The current model also leverages the power-coupling between the Hopf-oscillators with different individual frequencies, harnessing theoretical advancement of coupled Hopf-oscillator system dynamics and devising a complex feedforward network to approximate the empirical signal. The model is composed of two cascades of stages:-in the first phase of learning, where the oscillators learn the frequency components of the teaching (EEG or BOLD) signals, the phase difference between the oscillations elicited by the oscillators, and in the second stage, a complex feedforward network is used. More details of the network architecture are provided in later “Methods” and “Results”(in the subsection named “Model Summary”) sections. Here, we develop a novel mathematical framework of a large-scale brain network model,– (1) introducing a unified mathematical framework with a network of oscillators simultaneously capturing EEG-fMRI data, incorporating the structural information, (2) developing a trainable Hopf-oscillator network-based whole brain model, where each brain region or ROI is represented by a small cluster of limit cycle Hopf oscillators instead of the single oscillator for one ROI, (3) deciphering the relationship between the frequency components present in the EEG and fMRI from the trainable Hebbian learning embedded in the model architecture, (4) reconstructing the empirical Functional Connectivity (FC) and Functional Connectivity Dynamics (FCD) obtained by analyzing EEG signals with different frequency bands and BOLD signals from the model with higher accuracy along with approximating the modularity in brain’s functional organization, (5) investigating the impact of structural disconnection on frequently measured matrices to track functional organization and network dynamics of the human brain, i.e., FC, FCD, modularity, and integration for low-frequency fMRI and high-frequency EEG from a single mathematical framework.

The article is organized as follows: The next section, the “Methods” or section 2 deals with the intricacies of the model’s architecture and the methods for analysing it. The “Results,” delves into the modeling study’s outcomes. Later, the “Discussion” section offers a critical review of the model required to give a broader overview and compare it to earlier developed models. The supplementary document attached to the study incorporates additional supporting results and the methodology.

## Methods

### Database Used

The dataset used in the model was sourced from an openly available concurrent EEG-fMRI signals’ data along with structural connectivity dataset. The similar dataset was used in previous study on concurrent EEG-fMRI modelling (Schirner et al., 2018)[Link to the database-https://osf.io/mndt8/]. Appropriate permissions were obtained for the use of similar datasets in this model. Among fifteen subjects between 18 and 30 years old (including eight female subjects), the first nine indexed subjects’ data are used in this study. The dataset contains already preprocessed fMRI and EEG data. The repetition time (TR) of fMRI data is 1.9 seconds, and 3 minutes and 14 seconds of data have been used for simulation purposes (excluding the first five scans’ data). The pipeline for preprocessing the fMRI data has been given in the earlier mentioned simultaneous EEG-fMRI modelling work by Schirner et al. (Schirner et al., 2018). The EEG data was preprocessed by the BrainVision Analyzer toolbox and later sampled to 200 HZ and low pass filtered to 60 HZ with the EEGLAB toolbox (Delorme and Makeig, 2004). Note that the validation dataset is obtained from the open-source “EEG-fMRI-NODDI” dataset provided by Deligianni et al. (Deligianni et al., 2016) [Link to the database-https://osf.io/94c5t/]. The details of the preprocessing steps are given in the subsection S.1.3 of supplementary document online. In this article, we refer to the data provided by Schirner et al. (2017) as the “Berlin dataset”, and the dataset provided by Deligianni et al. (2016) as “EEG-fMRI-NODDI dataset”.

### Basic Mathematical Model

The phenomenological model proposed in the current study is based on a network of non-linear harmonic oscillators – the Hopf oscillators - and strives to accurately match or mimic both the haemodynamic and electro-physiological data across two distinguished temporal scales. The modelling approach leverages the property of perturbation of the limit cycle oscillator and characteristics of Hopf oscillators to adapt the frequency of external signals, rendering the opportunity to reproduce the hemodynamic and electrophysiological signals. (Buchli et al., 2006; Righetti et al., 2005, 2009). In our earlier model (Bandyopadhyay et al., 2023), each brain region was represented by a single Hopf oscillator; but in the current study each ROI is represented by a network of Hopf oscillators. Thus, presently, the whole brain dynamics is modelled by a network of networks. The network is trained using two stages of learning, which are discussed below.

### First Phase of Learning

The objective of the current model is to reproduce the EEG and BOLD signals simultaneously. We just mentioned that each ROI is represented by a network or a population of Hopf oscillators. This population is further divided into two clusters: the low frequency and the high frequency clusters. The low frequency oscillators (LFO) with the frequency range 0.01−0.09 Hz (intrinsic frequency as initialized) are designated to model the low frequency oscillations of the BOLD signal. Oscillators with intrinsic frequency as initialized with the range of 1.5 − 49 Hz representing the much faster EEG signals are called HFO (High-frequency oscillators). In Fig. 1 (a), both the clusters of oscillators (LFO and HFO) are shown at different levels according to their intrinsic frequencies. All the oscillators corresponding to a single ROI are connected in an all-to-all manner. There are three types of connections among the oscillators : intra–cluster, inter-cluster, and EEG-fMRI connection. Intra-cluster connections are all-to-all connections of oscillators inside each sub-cluster of same cluster (ROI), irrespective of HFO or LFO levels. In the current framework, we studied a minimal set of five oscillators that pose all-to-all connections. Inter-cluster connection refers to the connections between the oscillators of different clusters (ROI) but at the same level-either in LFO or HFO, and the EEG-fMRI connections refer to the connections between the LFO and HFO oscillators in the same clusters. The oscillators in the low frequency cluster are used to reconstruct fMRI signals while the high frequency oscillators are used to reconstruct the EEG signals.

**Fig. 1:**
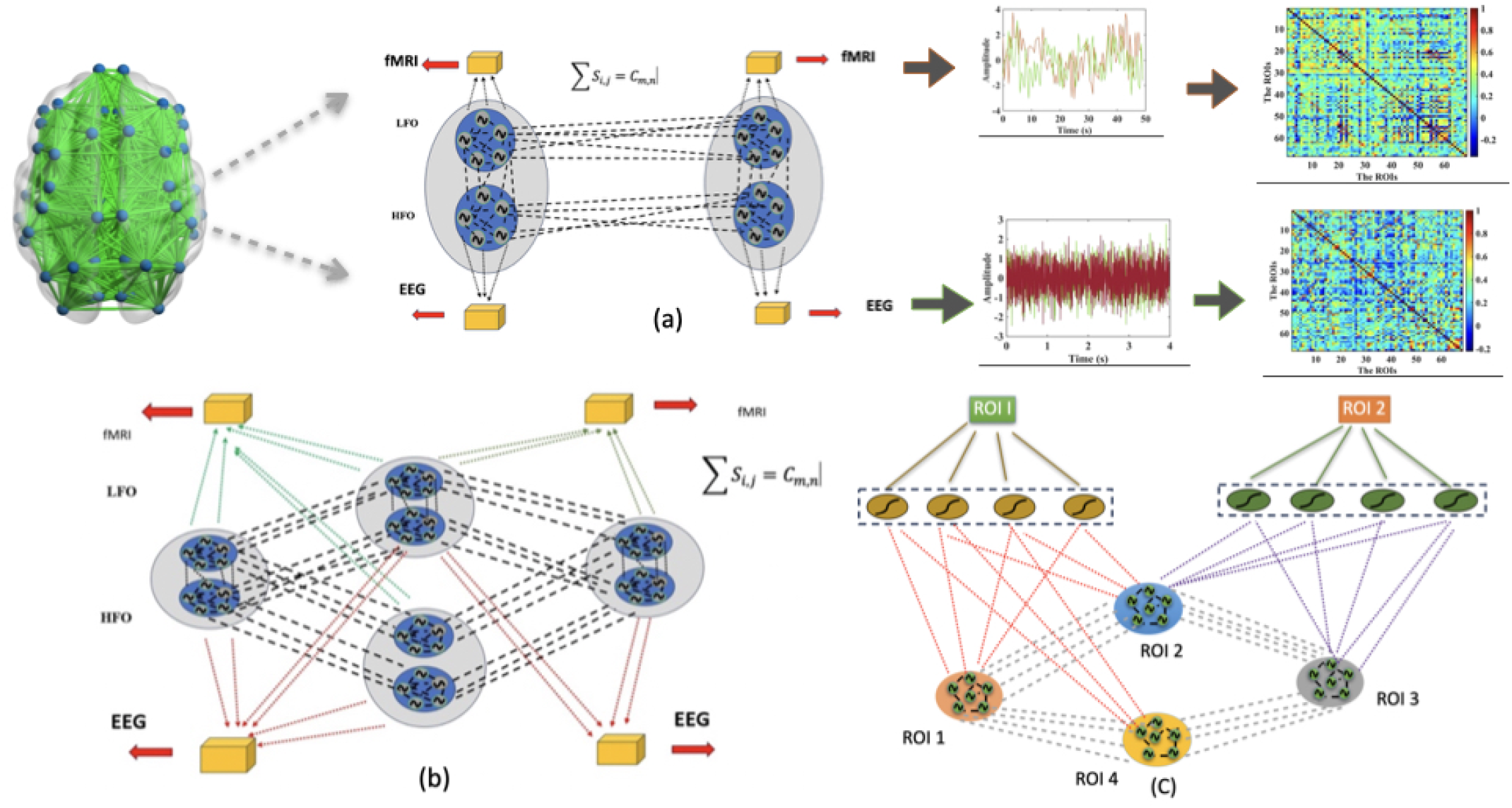
A schematic diagram of the mathematical model. Fig. (a) represents how LFOs and HFOs embedded in two ROIs take part in training algorithm and the connections in-between the ROIs remain constant, i.e. ∑ *_i,j_ S_i,j_* = *C_m,n_*. In Fig.(b) four interconnected ROIs participate (as an example) in first phase of learning. As mentioned in the text, the ROIs which are connected to any *i^th^* ROI take part in producing *i^th^* signal irrespective of EEG or fMRI signal. Fig. (c) shows the complex-valued feedforward network used to approximate the EEG signal.

Characteristic equation of a network of Hopf oscillators can be given by (Biswas et al., 2021; Bandyopadhyay et al., 2023) —

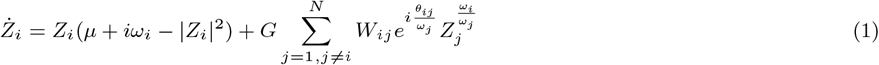

where, where, *Z* can described in both polar form (*r, ϕ*) and cartesian form (*x, y*). *µ* is the bifurcation parameter controlling the both the dynamics and amplitude of the Hopf oscillation. We take the Hopf oscillator in periodic oscillatory regime with *µ* = 1. The *ω_i_* represents the intrinsic frequency of the Hopf oscillator. For LFOs the *ω_i_* are initialized between 0.01 − 0.09 Hz (*ω_iLF O_* ∈ {0.01, 0.09} Hz), and for HFOs the *ω_i_* are randomly initialized between 1.5−49 Hz (*ω_iHF O_* ∈ {1.5, 49} Hz). Typically, the *W_ij_* refers to the adjacency matrix of the network of oscillators which represents the pattern and strength of the connections among the oscillators. Here, there are three different types of connections as described earlier, which can be referred as —intra–cluster (*W_ir,jr_*), where *i^th^* oscillator from *r^th^* cluster is connected to *j^th^* oscillator within cluster *r^th^* (*i^th^* and *j^th^* oscillator should be from either LFOs or HFOs); inter-cluster connections (*W_ir,js_*), where *i^th^* oscillator from *r^th^* cluster is connected to *j^th^* oscillator of *s^th^* cluster (*i^th^* and *j^th^* oscillator should be from either LFOs or HFOs), and inter EEG-fMRI connection (*W_if,je_*), where *i^th^* oscillator from *f^th^* cluster corresponding to LFOs connect to *j^th^* oscillator from *e^th^* cluster corresponding to HFOs (*i^th^* and *j^th^* oscillator should represent LFOs and HFOs respectively or vice-versa). Readers may refer to the schematic diagram given in the Fig. 4(b) for three different types of connections. The connection between a pair oscillators is a complex number and hence depicted by the amplitude, and the angle. The amplitude part shows the connectivity strength between the oscillators, and the angle is given by *θ* corresponds, broadly speaking, to time delay. Note that, the current form of coupling, as shown in the equation 1, is the novel “power coupling”, which is the revised complex coupling. It emerges as the unique coupling strategy that can tackle the well-known problem of coupling of two or more Hopf oscillators with different intrinsic frequencies. Such coupling shows that normalized phase difference 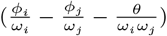 remains constant with time, and make the model more biologically plausible (Biswas et al., 2021). Our earlier work shows the usefulness of “power coupling” in modeling whole brain dynamics and yielded results comparable to the well known virtual brain model of Deco et al (Bandyopadhyay et al., 2023; Deco et al., 2017b). A closer look at equation 1 reveals that in power-coupling, the complex state of the oscillator is raised to the power given by the ratio of the intrinsic frequencies,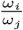. However, as the electrophysiological data and hemodynamic data, and *ω_iLF O_* and *ω_iHF_* have a large difference in frequency range, it’s imperative to restrict the ratio between *ω_iLF O_* and *ω_iHF O_* in a bounded manner, and to mitigate the problem, instead of linear scale, a logarithm scale is used. Interestingly, brain rhythm in logarithm scale showcases unique characteristics as revealed by Penttonen and Buzsaki (2003), like the frequency bands of varied brain rhythm form a geometric progression with constant ratio of 2.72 in a linear scale, which translates into an arithmetic progression in logarithm scale rendering the access of wide range of frequency bands of oscillations - from very slow (BOLD signal) to fast (EEG) frequency bands (Penttonen and Buzsáki, 2003; Buzsaki and Draguhn, 2004). Kim and Large has used such approach in their Gradient Frequency Neural Network (GrFNN) to understand the complex coupling between two or more Hopf oscillators (Kim and Large, 2015, 2019; Humphries et al., 2010).

Figure 1 (b) depicts the first phase of learning, where the four clusters represent different ROIs, with each of them having a set of intra- and inter-connected sub-clusters of LFOs and HFOs. In this stage of learning, three separate learning rules are employed to learn the intrinsic frequency, angle of the “power-coupling” and the forward connections to the output. The LFOs and HFOs are learnt respectively for BOLD signal and EEG signals (in Fig. 1 (b) the forward connections for oscillator to output node for EEG and fMRIs are identified by a distinct colour codes). The mathematical expression of the learning rule is given by (Bandyopadhyay et al., 2023; Biswas et al., 2021) —

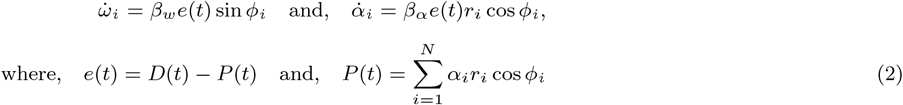

*β_ω_* and *β_α_* are the corresponding learning rates, *D*(*t*) is the learning/ teaching signals, and *e*(*t*), the error, is defined by the difference between simulated signal, *D*(*t*) and predicted signal, *P* (*t*). The *ω_iLF O_* and *ω_iHF O_* are trained with respective BOLD and EEG signals, and the lateral connections between the oscillators —the amplitude part is constrained by fixed value, and the angle is updated with a revised complex Hebbian learning rule as given in the following equation 3.

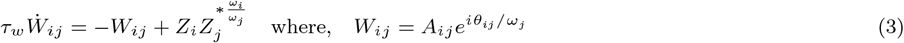

The amplitude of the intra-cluster connections as mentioned earlier can be represented by |*W_ir,jr_*|, which are fixed at 0.01, and for the aformentioned inter-cluster connections *W_ir,js_*, the amplitude is constrained such a way that the summation of them is equal to the actual fiber strength between two ROIs, i.e.∑ *_i,j_*|*W_ir,js_*| ∑ = *S_i,j_* = *C_m,n_* where, the *C_m,n_* is the connectivity strength between two ROIs. More about the parameter values are given in the parameter table Table S1 in the supplementary document.

Another important aspect of the model is that how the ROIs and associated cluster of oscillators are selected. To simulate the BOLD and EEG signals of the *r^th^* ROI concurrently, the number of oscillators, *n*, in *r^th^* cluster/ROI will connect to n number of oscillators in *s^th^* cluster if the *r^th^* ROI is structurally connected to *s^th^* ROI as deduced from the SC matrix. In later cases to decipher the neural basis of the hemodynamics signals, the amplitude of the connections between the two sets of oscillators with disparate intrinsic frequencies for HFOs, and LFOs, *W_if,je_* is trained with Hebbian rule, with bounds on the amplitude (|*W_if,je_*| ∈ {0.005, 0.05}) along with the the angle (*θ* as given in the aforementioned equation 1) between them. This has been discussed in the “concurrent EEG-fMRI data analysis” section again.

For simulation purpose, the timestep is taken to be 0.01 seconds, and the EEG signals and BOLD signals are respectively downsampled and upsampled to 100 Hz. Noted that, such adjustment of sampling frequency was adopted in previous studies also to make the simulation time-step match with the signals’ sample frequency (Schirner et al., 2018).

### Second phase of learning

After the first phase of learning, with intrinsic frequencies of the LFOs initialized along a logarithmic scale, it is possible to achieve a good approximation of the BOLD signal, but that is not the case with the EEG signals. As we have shown earlier, the complex-valued weighted feedforward network with oscillators enables to learn the amplitude of a signal (Bandyopadhyay et al., 2023), which is not possible with the first phase of learning alone. To capture the wide range of frequency patterns, at first the EEG signals are learnt with the first phase of learning to infer the frequency component and phase information with Hebbian learning, and later the oscillators with learnt frequency and with updated connection-information are trained with this hidden-layered feedforward network. Capability of such network, and the building block of it has been discussed at length in our previous studies where it is shown that the model accuracy is proportionally dependent on number of neurons in hidden layer and number of epochs (Bandyopadhyay et al., 2022). In the current framework we fixed the number of epochs at 10000, and number of hidden nodes at 20. The model’s performance is also dependent on the number of oscillators in each cluster. We set the minimal number of oscillators at five based on the performance of the model, and also provide the result for ten oscillators in each clusters to show the improvement in accuracy for the model to approximate each signal —be it BOLD or EEG signals. A schematic diagram of such a network is given in Fig. 1 (c). In the current framework, each ROI is represented by an ensemble of oscillators. We briefly discussed the mathematical rules for the complex feedforward network in the Supplementary Document online (refer to Subsection-S.1.1.2). Readers are requested to refer to our earlier works for the network building strategy and characteristics of this network (Bandyopadhyay et al., 2022, 2023).

### FC, FCD, and Graph Theoretical Measures

FC for the haemodynamic (fMRI) data is calculated by estimating the linear statistical dependency of the two ROIs in form of Pearson’s correlation coefficient. For estimating the FC for the electro-physiological data, at first the EEG data is band-passed for different frequency bands including delta (2 − 4 Hz), theta (4 − 8 Hz), alpha (8 − 13 Hz), beta (13 − 30 Hz), and gamma (*>* 30 Hz) frequency ranges. In the current study the Amplitude envelope correlation or AEC is estimated, which is widely used for computing the FC (Castaldo et al., 2023; Wirsich et al., 2021). For calculating the amplitude envelope, at first the analytical signal is evaluated with the Hilbert’s transform, and the amplitude value is taken. For across the frequency bands, AEC is calculated by taking Pearson’s correlation coefficient between estimated envelopes from each of two EEG signals from two ROIs, respectively(e.g. *i^th^* signal from *i^th^* ROI and *j^th^* signal from *j^th^* ROI, where, *i, j* ∈ {1, 68}). For capturing FCD, the time-series signals are segmented by sliding window technique with particular length and overlap or stride. For EEG data, the window length is fixed at 2 seconds with 50% overlap, and for BOLD signal it is 10 seconds with 40% overlap. For each cases different *N_τ_* numbers of FC can be found and a *N_τ_* × *N_τ_* FCD matrix can be deduced from estimating the Pearson’s correlation between two FC matrices at each window, *κ*, where *κ* ∈ {1*, N_τ_* }). Note that, for each element (say, *κ*(1),*κ*(2)) in the *N_τ_* × *N_τ_* FCD matrix denotes the correlation coefficient between upper or lower triangular elements of FC matrices, i.e. *corr*(*FC*_*κ*(1)_, *FC*_*κ*(2)_). This methodology peruses the earlier well established protocol followed in multiple studies (Castaldo et al., 2023; Deco et al., 2017b).

One of the better ways to understand the topological attributes of the graph is to evaluate how modular the graph is. To this end, we have taken the Louvain Modularity algorithm provided by Blondel et al. (Blondel et al., 2008), and given by the following equation-

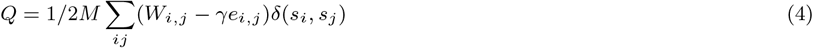

Here, *M* is the total number of edges in the network (*M* = 1*/*2 ∑ *_i,j_ W_i,j_*), and *W_i,j_* is the weighted connectivity/adjacency matrix of the network. *e_i,j_* can be obtained following the configuration model (*e_i,j_* = *k_i_k_j_/*2*M*), where *k_i_* and *k_j_* are the degrees of nodes i and j, respectively. *δ*(*s_i_, s_j_*) represents the Kronecker’s delta function showing if two nodes, i.e. *i^th^* and *j^th^* node are in the same modules or not. Note that *γ* is the resolution parameter controlling the number of partitions. The Q value represents the modular architecture of the network, i.e higher intramodular and lower intermodular connectivity than appearance by chance. However, such Louvain-like modularity estimation algorithm has an intrinsic (degeneracy) problem, which can be somewhat, negated by the multiple runs of the algorithm, which reveals each node’s affiliation to a module measured by the number of times a node is assigned to a particular community/ module (Lancichinetti and Fortunato, 2012; Good et al., 2010). Pursuing the earlier works, a matrix *D_i,j_*, referred to as a co-classification matrix or module allegiance matrix, is estimated representing the probability of the *i^th^* and *j^th^* nodes to be assigned in the same community for all the iterations (Lancichinetti and Fortunato, 2012; Good et al., 2010). The community structure formed from both the simulated and empirical EEG and fMRI data has been evaluated. Note that, in the current study, a total of nine subjects’ simultaneous EEG and fMRI data have been evaluated, and a “representative” functional connectivity matrix has been formed by taking the mean connectivity matrix of those nine subjects’ haemodynamic and electrophysiological data.

To compute dynamic modularity, instead of evaluating the modularity for each time-windowed functional connectivity, Muncha et al.(2010) provided a more elegant solution to this non-trivial problem by suggesting a Louvain-like modularity function taking into account the multilayer, multiplex network (Mucha et al., 2010). The revised quality function looks like-

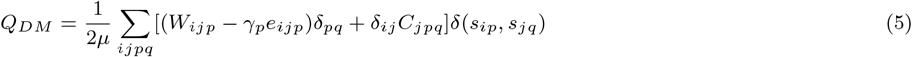

Here, *e_ijp_* estimates the configuration null model based on Newman-Girvan model for each window level, p and defined by,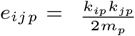, and 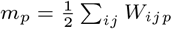. Note that the two variable-parameters in the model are - resolution parameter, which is kept constant as 1.1 (default value-1), and *C_jpq_* defines the connection strength between the layers of the multislice network. Interestingly, the equation holds the information of both the connections of a node in a single layer given by the first half of the equation (*W_ijp_* − *γ_p_e_ijp_*)*δ_pq_* defines the modularity for a single layer, where the second part of the equation given by the *δ_ij_ C_jpq_* couples the subsequent layers. *µ* is the normalization factor taking into account the total number of connections/edges in each layer and in-between layers, which can be described by 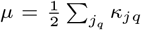, and *κ_jp_* = *k_jp_* + *C_jp_*. Here, *k_jp_* represents how a node j is connected to all nodes in a layer p, and how it is connected across layer in a multislice network is given by *C_jp_* or *C_jp_* ∑= Σ*_q_ C_jpq_*. As suggested by the original work, the *C_jpq_* is defined by *C_jpq_* ∈ {0*, ω*} based on the presence and absence of multislice network and the *ω* has been taken as 1 in current study. Updated version of GenLouvain algorithm is used in this regard, which is available in online repository (Jutla et al., 2011) [it’s available on following online repository-https://github.com/GenLouvain/GenLouvain].

### Concurrent EEG-fMRI data analysis

To decipher the intricate relationship between high frequency EEG signals and the slow frequency fMRI signals, two different approaches are adopted in the current study via., data-based approach wherein conclusions are drawn by analyzing the data, and the model-based approach where the model parameters, obtained by the training algorithm, reveal the mapping between neural correlations with different EEG frequency bands and the fMRI signals’ FC. For the first approach, AEC FCs for different EEG frequency bands have been evaluated. Similarly, the Pearson’s correlation based FC for BOLD data is also estimated for both simulated and empirical data of each participant. A grand average AEC FC for all the frequency bands and FC for fMRI signals is obtained designating a “representative” correlation structure or the functional connectivity map for human brain with two distinct time-scale and frequency ranges. Pursuing the earlier works, correlation between two functional connectivity maps have been obtained to find out the relationship between them (Wirsich et al., 2021). It is noteworthy that total nine subjects’ data have been taken from available fifteen subjects data from Berlin dataset to find the “grand average” as the earlier works show that at least seven or more than that number of subjects’ is required to get a stable, significant relationship (Wirsich et al., 2021).

However, as previously stated, the data-based analysis is marred with obscurity and a lack of standardization, while the current mathematical framework has the capacity to address this challenging issues. As stated earlier in the “Basic Mathematical Model” section, there are three separate kinds of connections (shown in (B) in Fig. 4(b)). The amplitudes of the three connections are fixed at a predefined value as stated in the “first phase of learning “ subsection under the “Basic Mathematical model”. However, at this end, among the three connections, the amplitudes of the inter EEG-fMRI connections or the |*W_if,je_*| are updated along with the intrinsic frequency of the oscillators and the angle of coupling (∠*W_if,je_*) of the “power coupling” connections between the oscillators. At the end of the training by the Hebbian learning as given in the equation 3, each connections are normalized and the amplitudes of the connections belonging to frequency of which oscillators are noted. It is important to note that the amplitude of the connections are restricted, i.e. |*W_if,je_*| ∈ {0.005, 0.05} for the Hebbian learning. Top one-percentile and five-percentile strongest connections between the LFOs and HFOs are taken and frequency range of the HFOs, which strongly couple between LFOs and HFOs representing slow frequency oscillations of fMRI and high frequency oscillations of EEG respectively, are estimated. In the current text, we focus on one-percentile strongest connections.

### Structural Prunning

The manner in which loss of structural information is manifested in terms of network dynamics can be understood with the help of in-silico perturbation of the whole brain connectome, where the connections between the ROIs are pruned either by global or focal lesion. Such structural pruning can give insights into the dynamical features of neurological disorders that arise due to structural disconnection. In our previous work, we showed that such virtual perturbation causes a significant anomaly in FC, FCD network (Bandyopadhyay et al., 2023). Here a systemic lesion in the brain connections is introduced by applying either an absolute threshold or a proportional threshold on the structural connectivity matrix. Absolute threshold refers to keeping the structural component from the structural connectivity matrix above a certain threshold value and resetting the other subthreshold elements of that connectivity matrix to 0. Here three threshold values have has been taken −0.5, 0.10, and 0.20. Similarly, for the proportional threshold, a certain percentile of stronger connections are kept intact keeping the comparatively weaker connections at 0.The top one, five and ten percentile stronger connections are kept intact in each case of perturbation. Note that, for each perturbation, we compare the functional information in terms of the functional connectivity, functional connectivity dynamic, community structure, and spatio-temporal integration.

### Spatio-temporal integration

To measure the spatio-temporal integration-segregation for haemodynamic data, a unique joint histogram of nodes’ graph theoretical parameters can be obtained to draw “cartographic profile”, a joint histogram of the participation coefficient and degree z score, revealing the individual node’s role in integration and segregation (Guimera and Nunes Amaral, 2005). Such a histogram profile carries vital information such as hub and non-hub nodes. For this procedure we followed the protocol from Shine et al, and Luppi et al. (Luppi et al., 2021; Shine et al., 2016). The protocol is as follows - (1) At first, the time-resolved functional connectivity has been calculated to map the functional connectivity dynamics and deduce graphical architecture of brain. (2) Graph-partitioning Modularity algorithm has been adopted with signed version (equation 4 showcases the unsigned version) to take into account for the negative connections, which can be presented by the following equation-

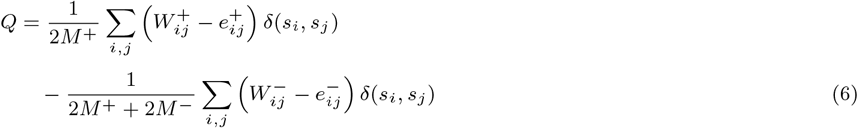

Here, *W_i,j_* refers to the weighted connection between node i and j; *M* refers to the total weight of the network, and the *δ* function estimates 1 when the two nodes *i*, and *j* are in the same module, and 0 otherwise. Note that, both positive and negative connections contribute for modularity evaluation represented by the sign in the superscript, and there is a asymmetric scaling of the signed weights by rendering more importance to the positive weights. This process has been done for multiple iterations (500 times) to find a consensus partition pursuing the earlier developed algorithm. Note that, for each step, the threshold for agreement matrix is set at 0.40. (3) After obtaining the partitions, the weighted and signed version of participation coefficient (*B_iT_*) and within-module degree z-score (*W_iT_*) has been evaluated to understand a node’s role in inter-module and intra-module connectivity resulting a cartographic profile representing brain’s functional integration and segregation. Weighted and signed version of participation coefficient (*B_iT_*) is given by the following equation (Fornito et al., 2016) -

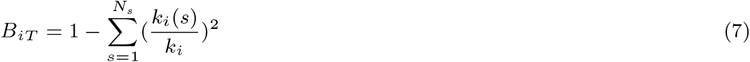

where *N_s_* is the number of modules in a network, *k_i_*(*s*) refers to the number of connections from node *i* to nodes in module *s, k_i_* refers to the sum of strengths of all connections. Note that, the *B_iT_* refers to the participation coefficient at each temporal window, T. The other parameter, within-module degree z-score (*W_iT_*) reveals the node’s functional role in intra-modular connectivity, given by (Fornito et al., 2016)-

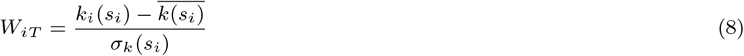

Here, *k_i_*(*s_i_*) refers to the within-module degree of node i within module *s_i_*, i.e. connections of node i with all other nodes, 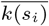, and *σ_k_*(*s_i_*) indicates the mean within-module degree of all the nodes in module *s_i_*, and standard deviation of all the *k_i_*(*s_i_*) values. (4) A cartographic profile for each temporal window is created by taking the joint-histogram of within and between module’s connectivity with both *W_iT_* and *B_iT_* parameters (the code developed by Shine et al. is used-https://github.com/macshine/integration/). An unsupervised clustering algorithm, k-means clustering (k=2) has been employed to distinguish predominantly integrated and predominantly segregated state by assigning the network derived from the temporal window having higher mean participation coefficient value as integrated state, and the other one segregated state. The k-means algorithm has been employed 5000 times to negate the initialization problem. This algorithm is repeated 5000 times with random re-initialization with pearson correlation to mark the distance metric in MATLAB’s K-Means implementation. The similar procedure is applied for both empirical and simulated BOLD signals. The proportion of “predominantly” integrated states emerging from the analysis is denoted by Integrated State of Occurrence Rate (ISOR). This methodology is also employed to estimate the integration state of the brain by evaluating ISOR for the two aforementioned pruning conditions of structural connectivity matrix, viz., absolute threshold and proportional threshold.

For calculating the integration level of the electrophysiological signal, an earlier established Nested-Spectral clustering/partitioning (NSP) algorithm has been adopted, which stems from the idea of spectral graph theory, referring a graph that can be partitioned/ clustered based on the eigen-decomposition of its adjacency/Laplacian matrix associated with it. The procedure, adopted from earlier work, and pursued here is as follows (Zhou et al., 2023; Wang et al., 2022) - (1) For each time-resolved FC, the symmetric FC can be written with its eigenvalues and corresponding eigenvectors, i.e. *C_t_* = *X*Λ*X^T^*, where *C_T_* is the functional connectivity matrix at each time window, *T*. The eigenvalues are arranged in decreasing order with corresponding eigenvectors. NSP defines clusters based on the eigenvectors in a hierarchical fashion and subsequent partitioning, resulting in different levels/orders of partitioning. For *i^th^* level the integration component can be quantified by (Wang et al., 2022; Zhou et al., 2023) –

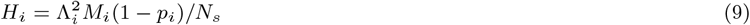

where, *p_i_* is the correction factor given by,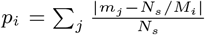. *M_i_* is the number of modules in the *i^th^* level of partitioning, *m_j_*being the size of *j^th^* module in each order of partitioning, and *N_s_* is the number of brain regions in the Desikan-Killiany atlas. For global integration, only the first level of partitioning has been considered, and it can be given by the following equation-

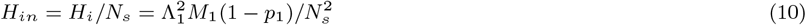

To evaluate the temporal fluctuations of the integration level, the integration level *H_in_* is computed across the temporal scale, which is given by the following equation-

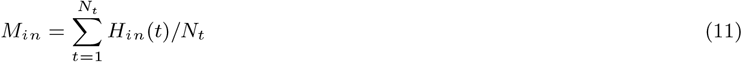

Note that, in the current study, only the integration level for both modalities is discussed as we hypothesize that a human brain can have only two states-integration and segregation; so both are complimentary to each other and vary in an inversely proportional fashion in quantity.

## Results

### Model Summary

The current work proposes a mechanistic oscillatory neural network model that can concurrently capture neuroimaging and electro-physiological data. In the current framework, each ROI (region of interest) is represented by a cluster of oscillators. Furthermore, this cluster is subdivided into two subclusters. In one of these clusters, the oscillators are low frequency oscillators, (*LFO*), with the frequency band corresponding to that of BOLD signal (0.01 − 0.09 Hz), whereas in the other cluster, the oscillators are high frequency of oscillators (*HFO*) with the frequency band corresponding to that of Electroencephalogram (EEG) (refer to the Fig. 2(a) (IV)). In the context of multi-modal simulation, specifically the simultaneous EEG-fMRI modeling in conjunction with empirical Diffusion Tensor Imaging (DTI) data, the network is designed to incorporate structural connectivity information. This ensures that the magnitude of coupling between oscillators within the same layer (SFO or HFO) from two distinct ROIs collectively reflects the structural connectivity between the ROIs. Moreover, these oscillators inside each sub-cluster are connected in an all-to-all fashion, and oscillators with disparate intrinsic frequencies (LFOs and HFOs) inside each cluster are also connected in all-to-all fashion (see the Fig. 2(a) (IV)).

**Fig. 2:**
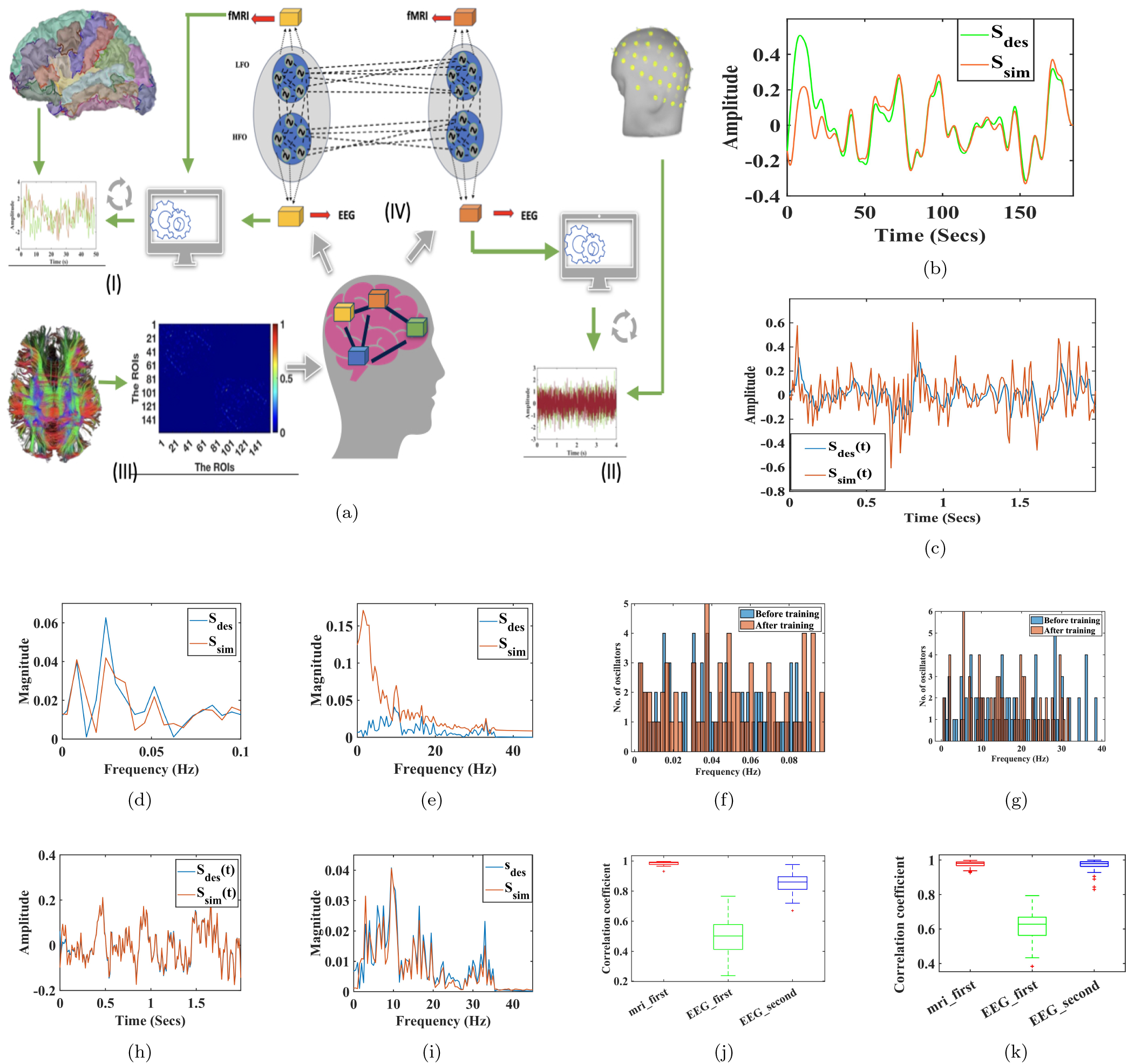
Reconstruction of concurrent EEG-fMRI data. Fig.(a) provides the schematic diagram of the model. Figs. (b) and (c) individually represent the model’s efficacy in capturing the BOLD and EEG signal after first phase of learning. Noteworthy, the *S_des_*(*t*) refers to the empirical signal, and the simulated signal is given by *S_sim_*(*t*). Figs. (d) and (e) reveal the models’ capacity to capture the spectral property by the first phase of learning itself for BOLD and EEG signals, respectively. Figs. (f) and (g) illustrate how the frequency of the oscillators is learnt by the first phase of learning for both *LFOs* and *HFOs*. Figs. (h) and (i) show the model’s effectiveness in capturing the time-domain and the frequency-domain data of EEG after the second phase of learning. Note that simulation data of only 2 seconds has been shown for EEG data. Figs. (j) and (k) assess the correlation coefficients between empirical and simulated signals with different numbers of oscillators (ten and twenty oscillators in each ROI individually).

Note that the limit cycle Hopf oscillator represents each LFO and HFO with their intrinsic frequencies. Several different types of coupling strategies have been in the literature, like-real-valued coupling and complex coupling; however, they are unable to maintain the stable phase difference or phase synchronization unless their intrinsic frequencies are almost identical. Hence, such coupling approaches are not reasonable and biologically plausible for a modelling framework for simultaneous EEG and fMRI data. In contrast, the “power-coupling”, as proposed in earlier works, is able to mitigate this problem by maintaining a normalized phase difference between the oscillators having different individual intrinsic frequencies (Biswas et al., 2021; Bandyopadhyay et al., 2023). A more comprehensive description of the methodologies used in this study is provided in the “Basic Mathematical Model” section. Noteworthy, the two subsets of oscillators in different layers (HFOs and LFOs in a ROI) are connected by an all to all “power-coupling’, where the connection strength between two oscillators or amplitude of power coupling corresponding to different frequency ranges are updated by a revised complex Hebbian learning rule in a restricted bound, enhancing the model’s effectiveness in elucidating the intricate relationship between low-frequency fMRI signals and high-frequency EEG signals.

The proposed model has two stages of learning: - in the first stage, the frequency of the oscillators and the lateral connections between oscillators, along with the forward connections to the output node, are updated. In the second stage, a feedforward network with the complex valued weights has been introduced to capture the empirical electrophysiological signal. This framework enables a nuanced and meticulous examination of neural dynamics, leveraging the distinct temporal characteristics captured by BOLD and EEG measurements. In later part a succinct discussion about the modelling simulation results have been illustrated.

### Reconstruction of Simultaneous EEG and BOLD signal

As outlined in the previous section and detailed in the “Basic Mathematical Model”, model simulations were done with the objective of reconstructing EEG and fMRI data simultaneously. In the current model, each ROI is represented by a cluster, which contains two sub-clusters of oscillators, corresponding to higher (HFO) and lower (LFO) frequency bands. These HFO oscillators and LFO oscillators are connected in an all to all fashion, referring each LFO oscillator in a subcluster of a ROI being connected to all the HFOs of that ROI and vice-versa. The minimal number of oscillators in each of the subcluster is set as 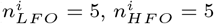, denoting number of LFOs and HFOs in each subcluster of *i^th^* ROI or *n^i^*=10, referring total number of oscillators in *i^th^* ROI), and the effect of taking more than five oscillators i.e. ten oscillators 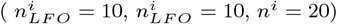 on simultaneous EEG and fMRI data simulation is also performed. A total of 194 seconds (3 minutes and 14 seconds) of recording of EEG and fMRI data is used for simulation. The present model is a personalized whole-brain model, in that it incorporates individual structural connectivity, EEG source activity data or EEG data collected from electrode projected to the cortical parcellation or Desikan-Killiany atlas (Desikan et al., 2006) and the fMRI data extracted from each ROI from the similar atlas for simulation of each individual. Details of the mathematical model and its intricacies are provided in detail in the “Methods” section and in the supplementary. In Fig. 2, the simulated outcome of concurrent EEG-fMRI or multimodal modelling framework corresponding to the both - first and second phase of learning is presented. The time and frequency domain signal analysed in Fig. 2 is from ROI indexed 1 from third subject’s data from the Berlin database (Schirner et al., 2018).[Link to the database-https://osf.io/mndt8/].

### Simulated outcome of the first-phase of learning

Figure 2(b) reveals that the current modelling approach approximately reconstructs the BOLD signal, and it can be observed from a time-series signal as well as frequency domain power spectrum as shown in the Fig. 2(d). Frequencies learnt from the model by the LFO oscillators are revealed in Fig. 2(f). The precision of the model to capture each of the BOLD signals from individual ROI is given in Figs. 2(j) and 2(k) for both five and ten oscillators in each subcluster, respectively 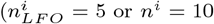 and 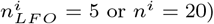. It reveals that the current mechanistic model can approximate the BOLD signal in the first phase of learning without even the second phase of the learning procedure. The tables are given in supplementary section (refer to Tables S3 and S4) illustrating the correlation value between simulated and empirical signal (here BOLD) also support this claim, where it is found that with five oscillators in each sub-cluster, 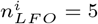 or *n^i^* = 10 the model can approximate the BOLD signal and marginal increase in the model’s accuracy is observed with ten oscillators in each sub-cluster or 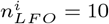 or *n^i^* = 20.

Efficacy of the first-phase learning in approximating the electrophysiological data can be obtained when observing the EEG simulated data (both in time and frequency-domain data) as depicted in Figs. 2(c), 2(e), and it’s important to note that the current modelling approach is unable to approximate the EEG empirical data in the first phase of learning alone. However, in the case of electrophysiological or EEG data, the oscillators learn the important frequency components of the pertaining EEG signals with the frequency learning algorithm, as shown in the Fig. 2(g). Closer observation will reveal that during HFO oscillators’ learning, most of the oscillators reach a narrow band of frequencies, primarily *α* and *β* bands. Interestingly, in multiple earlier works, it has been shown that resting state EEG is mostly occupied by *α* and *β* band regions, with strongest cortical correlation in that particular frequency range (Hipp et al., 2012; Hipp and Siegel, 2015).

### Simulated outcome of the second-phase of learning

As revealed in the earlier subsection, due to the inability of the first phase of learning to accurately capture the electrophysiological data, a complex-valued feedforward network is inserted between the oscillatory network and the output layer (refer to Fig. 1(c) in the “Basic Mathematical Model” section) in the second stage of learning, to achieve a more accurate fit of the empirical EEG signal. Insertion of a sizable hidden layer between the output of the oscillatory network and the final output, increases the approximation capability of the entire model. Interestingly, a similar approach, i.e., the second phase of learning, has been taken earlier to simulate much longer BOLD signal data, where a single oscillator has been assumed to represent a ROI (Bandyopadhyay et al., 2023). The model output approximates the EEG signal-both in the temporal and spectral domains accurately (Figs. 2(h) and 2(i)). Figures 2(j) and 2(k) demonstrate the predictive power of the model in capturing each ROI’s BOLD and EEG signal simultaneously, where they present the correlation coefficients among the empirical BOLD signal and the simulated BOLD signal, as well as between the empirical EEG signal and the simulated EEG signal after both the first and second phases of learning. Figures 2(j) and 2(k) show simulated results for 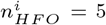 *n^i^* = 10 and 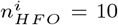 or *n^i^* = 20, respectively. These outcomes illustrate that with an increase in the number of oscillators in each cluster or ROIs, the model’s accuracy in capturing the EEG signal is increased both after the first and second phases of learning, where it shows only a marginal increase in the case of the BOLD signal. The similar pattern is reinforced by the observations given in Tables S3 and S4 in the supplimentary section (Refer to S2.1 subsection), where model’s simulated outcome (in term of correlation between simulated and empirical signal) for electrophysiological data for a single channel is provided. Also, it reveals that the second phase of learning has the capability to learn the amplitude of the EEG signals, which can be observed in both Figs. 2(j) and 2(k). There is a significant increase in the predictive power or correlation between the simulated and empirical EEG signals for each ROI after the second phase of learning, compared to the first phase. This improvement is also reflected in Tables S3 and S4, which show that the second phase of learning enhances the model’s ability to simulate the EEG signal of a single ROI. The next section will provide a further analysis involving model’s ability to reconstruct the functional connectivity map of the entire brain.

### Functional Connectivity and Functional Connectivity dynamics

Functional connectivity (FC) refers to the statistical relationship between the empirical time-series signals extracted from different ROIs, which informs about the functional organization and the hierarchical information processing in the human brain. In the case of a slow brain rhythm like the fMRI data, the functional relationship between two brain areas can be measured by taking Pearson’s correlation between the time series associated with the two brain regions. FC estimation for electro-physiological data can be obtained by either a time-resolved (Pearson’s correlation, mutual information, Granger’s causality) or a phase-based connectivity pattern (phase lock value (PLV), phase slope index (PSI), weighted phase lock index (wPLI)) (Sadaghiani et al., 2022; Abeysuriya et al., 2018). However, capturing electro-physiological data with non-invasive electrodes may lead to volume conduction artifacts due to erroneous localization on a larger spatial scale (Bastos and Schoffelen, 2016). This problem has been addressed using advanced source reconstruction algorithms that project electro-physiological data onto the whole brain parcellation obtained from neuroimaging data (Here Desikan-Killiany atlas has been taken).

One of the major benchmarks to validate a whole brain model is to check how the model can capture the functional connectivity and functional connectivity dynamics (Castaldo et al., 2023; Deco et al., 2017b; Cabral et al., 2017). In this section, we demonstrate the model’s accuracy by comparing the functional connectivity derived from empirical electrophysiological and neuroimaging (fMRI) data with the simulated FC, including functional connectivity dynamics (FCD), which captures the temporal variations of functional connectivity.

Our model, built upon a network of oscillators, successfully captures FC matrices emerging from amplitude envelope correlation (AEC), facilitating a detailed comparative analysis of FC and FCD estimated from simulated and empirical electrophysiological data (Fig. 3). Following previous research, the FC on EEG data is assessed in a frequency-resolved manner, wherein the raw time series data is filtered with a band-pass filter, including delta (2 − 4 Hz), theta (4 − 8 Hz), alpha (8 − 13 Hz), beta (13 − 30 Hz), and gamma (*>* 30 Hz) frequency ranges. However, only the connectivity pattern for the alpha band is presented here as the alpha band connectivity is found to be more reliable and consistent across subjects (Wirsich et al., 2021; Colclough et al., 2016; Marquetand et al., 2019; Nakuci et al., 2023). In Figs. 3(a) and 3(b), it is shown that the simulated AEC FC (Amplitude envelope correlation-based functional connectivity) approximately replicates the empirical one. The accuracy or predictive power of the model over the key EEG bands and across the subjects is shown in Fig. 3(e). Noteworthy, the predictive power implies the correlation coefficient between the simulated and empirical FC (noticeably, as both are symmetric in nature, only the upper or lower-triangular elements are taken for evaluation). On the other hand, the comparison between the two functional connectivity dynamic matrices can be estimated by calculating the distribution of correlation values from FCD matrices in terms of a non-parametric measure of distance between two distributions, namely Kolmogorov–Smirnov (*KS_distance_*) distance. Detailed discussion on the methodology adopted for FCD measurement is given in the “FC, FCD and graph theoretical measure” subsection in “Methods”. The procedure followed for this purpose is adopted from an earlier work, which has a different mathematical model to capture the simultaneously recorded MEG-fMRI data (Castaldo et al., 2023). Figure 3(f) reveals the model’s efficacy in approximating the empirical FCD matrix across all the subjects, as well as for different bands of EEG, as demonstrated by low *KS_distance_* distance, which refers to the distance between the distribution of empirical and simulated FCD values. An example of such distribution is shown in Figs. 3(i) and 3(j) demonstrating the FCD distribution between simulated and empirical for EEG-*α* and BOLD data. As the efficacy of the model is analyzed for EEG data, a similar analysis of the simulated BOLD signals are done for all the subjects. The Fig. 3(g) demonstrates that the predictive power of the model for BOLD data lies between 0.97 to 0.99, which convincingly outperforms the existing whole brain models that capture the simultaneous EEG and BOLD signals. Figure 3(h) denotes the Kolmogorov-Smirnov, *KS_distance_* for BOLD data for all the subjects, which supports the superiority of the model. However, as discussed before, the correct way of analyzing the EEG FC is still debated as the underlying artifacts arising from volume conduction. Interestingly, the current model is not susceptible to that problem as the data is collected from the DTI-parcellated ROIs.

**Fig. 3:**
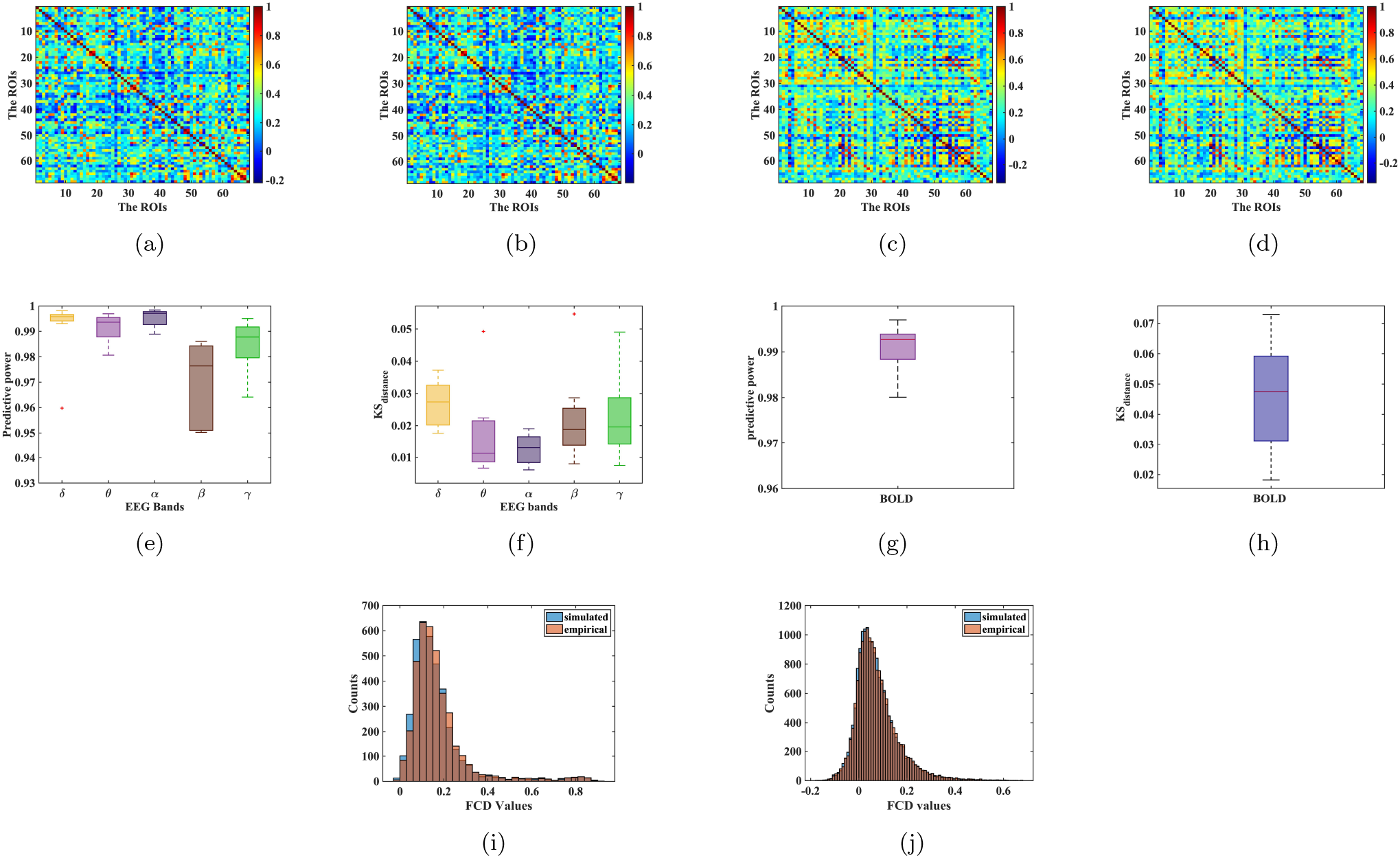
Comparison of functional connectivity map from both simulated and experimental data. Figs. (a) and (b) individually represent the FC (simulated and empirical) approximation of *α* frequency bands of EEG data with our model (only FC for *α* frequency band is shown here). Figs. (c) and (d) individually illustrate the simulated and experimental functional connectivity for BOLD signal, and Figs.(e) and (f) present the predictive power and dynamic functional connectivity represented by the distribution of FCD values for the nine participants of the test dataset for simulating all the key frequency bands of EEG with concurrent EEG-fMRI model. Figs. (g) and (h) represent predictive power and dynamic functional connectivity for BOLD signals. Figs. (i) and (j) reveal the histogram of the FCD for simulated and empirical data for EEG-*α* and BOLD data, respectively.

### Analysis of Concurrent EEG-fMRI data

Recent advancements in neuroimaging techniques enable us to decipher the functional connectivity map between the fMRI correlation and neuronal correlation estimated by frequency resolved FC evaluated from EEG data and FC estimated from BOLD data to elucidate the neural basis of hemodynamic signal, which remains somewhat ambiguous. FC matrices for both modalities are evaluated by analyzing the statistical dependency between different ROIs, which may or may not be structurally connected, for two different modalities operating on very different timescales. Multiple earlier works elucidate a significant relationship between the two modalities (Hipp and Siegel, 2015; Wirsich et al., 2021, 2020). Usually, this relationship is evaluated by comparing the correlation between AEC (Amplitude Envelope correlation) FC for EEG and Pearson’s correlation-based connectivity for fMRI. Adopting a similar methodology, we found that neural correlation for different frequency spectra is significantly coupled with the correlation matrix of hemodynamic signals of slow oscillation. Fig. 4(a) illustrates that AEC FC for EEG is correlated with Pearson’s correlation-based functional connectivity of fMRI signals for both simulated and empirical data, and noticeably, the overlap between fMRI correlation and *α*-and *β*-band AEC FC is higher than any other frequency bands, with the *β* band exhibiting the highest similarity to fMRI correlation structure. These findings align with earlier studies indicating a significant relationship between EEG data in the 8 − 32 Hz frequency range and fMRI data, particularly in multi-modal non-invasive approaches such as concurrent *EEG* − *f MRI* (Hipp and Siegel, 2015; Wirsich et al., 2021). It is important to note that, in addition to AEC, other connectivity measures, such as imaginary coherence, have been considered in earlier studies. However, we exclude this measure because it is more prone to noise-induced artifacts compared to AEC FC (Glomb et al., 2020).

Nonetheless, there are multiple caveats in this data-based analysis because of its problem with reproducibility, lack of standard protocol (preprocessing steps involved in acquiring the data) and a predefined statistical analysis of the correspondence between two different correlation matrices. Hipp et al. showed that SNR (Signal to noise ratio) corrected, and non-coincidental correlation (taking into account the dimensionality of the data) structure suggest that this relation is not confined to only *α* and *β* frequency range (Hipp and Siegel, 2015). Similarly, electrophysiological data obtained by invasive methods suggest that the strongest coupling exists between *γ* frequency band and correlation structure from haemodynamic data (Schölvinck et al., 2010; Mukamel et al., 2005). However, non-invasive studies suggest that the strongest correlation is prevalent between the correlation structure of *α, β* frequency range and fMRI data (Wirsich et al., 2021; Hipp and Siegel, 2015). The current study adopts the approach of the earlier work of Wirsich et al. (Wirsich et al., 2021).

To address the ongoing debate and dispute in this regard, we propose a mathematical model that might mitigate the problem. See subsection “Concurrent EEG fMRI data analysis” in “Methods” for a detailed description of the proposed methodology. In short, we adopted amplitude learning of the prevalent connections between LFOs and HFOs as shown in Fig. 4(b). The *LFO* − *HFO* connections, reflecting the inter EEG-fMRI connections (*W_if,je_*) in a ROI, are trained with the Hebbian learning rules (given in the eqn. 3), where *i^th^* oscillator from *f^th^* subcluster corresponding to LFOs connects to *j^th^* oscillator from *e^th^* sub-cluster corresponding to HFOs in a single ROI. At the end of the training, the top one percentile strongest connections and their corresponding individual frequency of HFO oscillators are noted. It is shown that the amplitude of the strongest coupling exists between *α* and *β* frequency ranges. Figure 4(b) illustrates a schematic diagram of the training procedure adopted, and the inset box in the Fig. shows that the updated weights are also normalized before extracting the statistical inferences out of it (mathematically,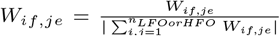). It is noteworthy that the update of the amplitude of the inter-connections between HFOs and LFOs does not compromise the precision or the model’s accuracy. Figure 4(c) showcases the schematic diagram of the connectivity pattern as emerged after the training iterations, where top 1% connectivity strength and the corresponding HFOs having stronger connections with the LFOs indicating the pattern of connectivity between low frequency oscillations and key frequency band patterns of EEG, are identified. On an average, how many LFOs are strongly connected (top 1%) to which frequency ranges of HFO for a single ROI are deduced during the simulation of all the ROIs and indicated in this figure. It shows that on an average, higher number of LFOs and HFOs pertaining to *α* and *β* are connected strongly. Figure 4(d) is a histogram plot of those top 1% HFOs for all the ROI’s simultaneous EEG-fMRI simulation and their frequency patterns, which are binned with 0.5 Hz width. It has been found that between the frequency range of 13 − 30 Hz, a typically *β*− band frequency range has the highest number of HFO oscillators having stronger connections with the LFOs.

**Fig. 4:**
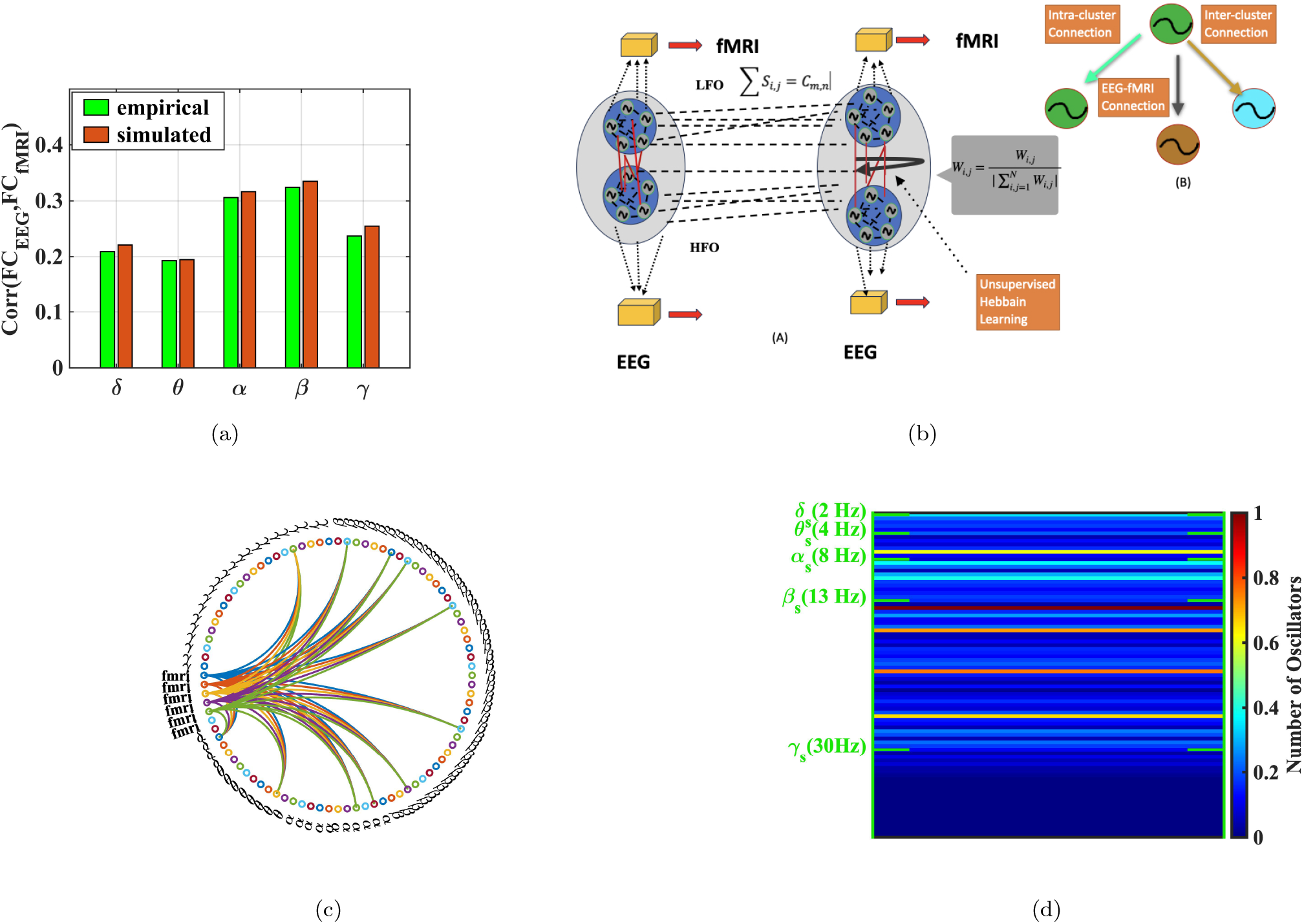
Inference from modelling and EEG-fMRI simultaneous data analysis. Fig. (a) reveals the correlation between AEC FC and Pearson-correlation based FC. Fig.(b) demonstrates a schematic diagram of the advanced version of the model, where the Hebbian learning-based training algorithm was optimized to train the amplitude of the connections between low (*LFO*) and high (*HFO*) frequency oscillators to deduce the relationship between electro-physiological and haemodynamic signals. Fig. (c) shows a schematic diagram of the connectivity between low-frequency *LFO* and high-frequency *HFO* oscillators, where the circular graph illustrates the connectivity pattern emerged for an individual ROI (on average) after the complex Hebbian learning. Fig. (d) reveals the connectivity strength of each *HFOs* with the *LFOs*. This is a histogram plot revealing intrinsic frequencies of *HFOs* adhere to the top 1% connectivity strength. The range of 0 − 1 represents the ratio of available number of oscillators of that frequency-bin (bin-width 0.5 Hz) to the total available number of oscillators. Note that only the starting frequency of each frequency band is indicated in the figure.

### Graph Theoretical measures (modularity) analysis

Modularity is one of the key attributes of graph theoretical measures of brain networks, deciphering how this network segregated in different communities manifests seamless functional information processing in brain. Such measures have been routinely used to explain the emergence of distinct modules from the functional connectivity analysis (Alexander-Bloch et al., 2010). To this end, modularity is evaluated from the network emerging from FC and FCD. Interestingly, modularity calculated from both time-resolved and without-time-resolved data provides information about the integration and segregation of the network (Bassett et al., 2015; Finc et al., 2020). Both Figs. 5(a) and 5(b) demonstrate the modularity value of the FC network derived from both simulated and empirical data, as well as the modularity estimation from a random network providing the null distribution of modularity from 100 rewired networks. Following previous work a normalized modularity value is calculated and shown in Fig. 5 (Finc et al., 2020). Normalized modularity refers to the ratio between modularity values estimated from the empirical data and mean-modularity value form the randomized networks. Interestingly, the results also support an earlier empirical study on simultaneous EEG-fMRI data analysis, where it was shown that the modularity value of the network generated from BOLD signals is higher than that of the network generated from various frequency bands of electro-physiological signals (Ayres-Ribeiro et al., 2023). Such normalized modularity calculation has been proved to be useful tool to understand the cognitive load for different training experiences and task conditions (Finc et al., 2020). Dynamic modularity is calculated for both simulated and empirical data for different bands of EEG and fMRI data, and the simultaneous EEG-fMRI model almost captures the modularity value estimation from empirical data obtained from both modalities. Noteworthy, such a modularity estimation algorithm is prone to algorithmic degeneracy due to modularity maximization algorithms, like-the Louvain algorithm, resulting in inconsistent modular affiliation of an individual node with almost similar modularity score for multiple iterations of the algorithm, and the problem can be mitigated by evaluating module allegiance matrix or co-classification matrix, providing a representative/consensus community structure (Good et al., 2010; Lancichinetti and Fortunato, 2012). A detailed description of the methodology is given in the subsection “graph theoretical measure” in “Methods” section. Figs. 5(e) and 5(f) represent the co-classification matrix for simulated and empirical BOLD signals individually. On the other hand, Figs. 5(g) and 5(h) reveal the co-classification matrix for simulated and empirical alpha EEG signals individually. As the time-windowed FC matrix fluctuates over time, and the modularity value estimated from individual time windows allows us to track the variance of modularity over time(Fukushima et al., 2018; Fornito et al., 2016). Note that, Muncha et al (2010) provided a unique way to understand such time-dependent multi-layer networks (Mucha et al., 2010). The corresponding methodology has been described in earlier in “Methods” section. Figs. 5(c) and 5(d) refer to the modularity value from the multi-layer network derived from windowed empirical and simulated data, and a comparison is done with a null model network derived by randomizing each time-windowed network. However, there are multiple ways to show the robustness of modularity estimation (Besset et al. 2013), which, however, is out of the scope of the current study (Bassett et al., 2013).

**Fig. 5:**
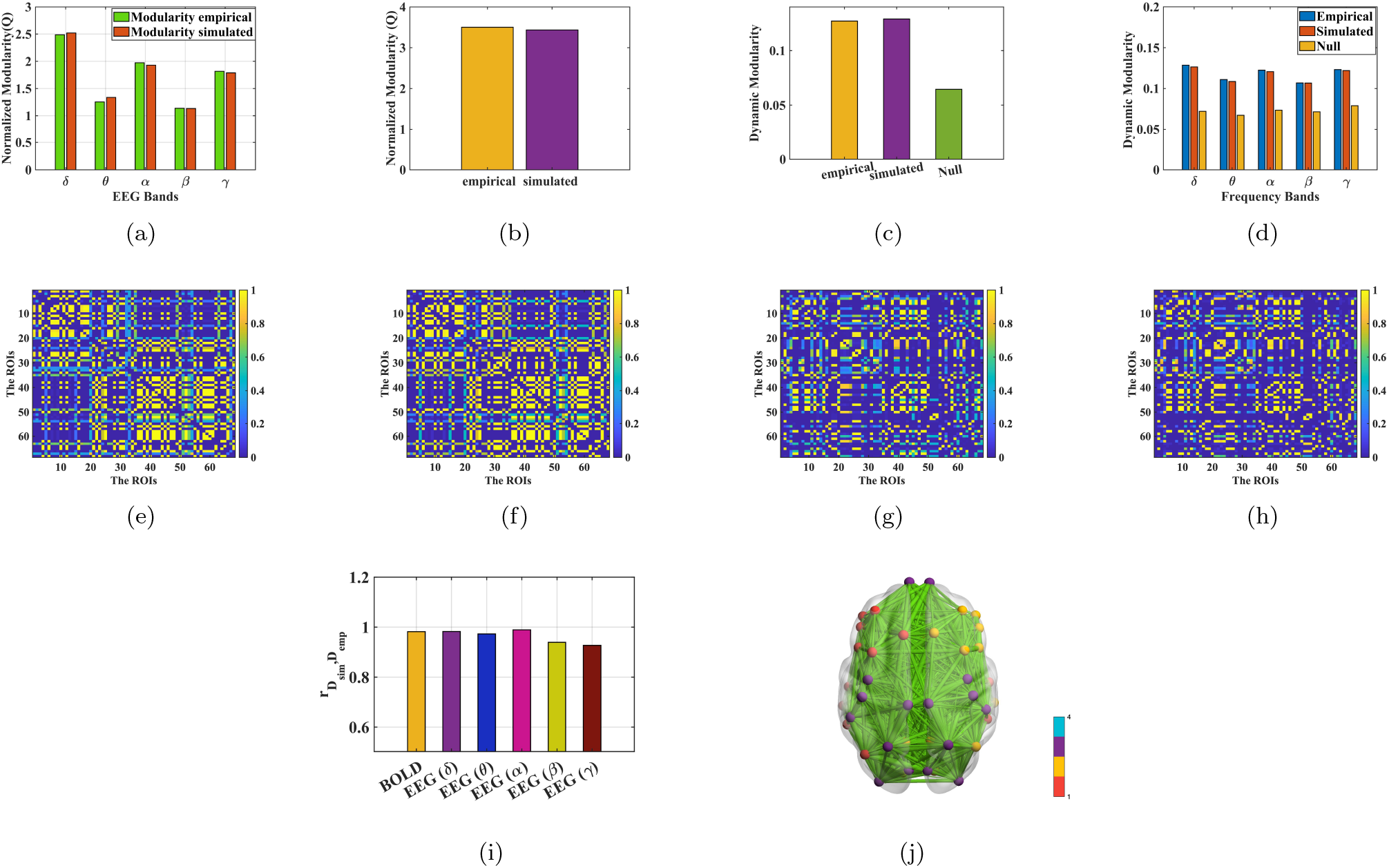
Graph theoretical measurement for simulated and empirical EEG and fMRI data. Figs. (a) and (b) represent the normalized modularity value for the network emerging from different frequency bands of EEG and the BOLD FCs. Figs. (c) and (d) demonstrate the dynamic modularity from multiplex network deduced from the time-resolved/windowed FCs by sliding window procedure for both fMRI and EEG data. Figs.(e) and (f) refer to the community structure estimated from simulated and empirical EEG-*α*. Figs. (g), (h) demonstrate the community structure for both empirical and simulated *f MRI* data. Fig. (i) compares the representative “community structure” estimated from simulated and empirical data for both different frequency bands of Electro-physiological and haemodynamic signal. Fig. (j) represents the schematic diagram of the graphical architecture of the brain, having modular architecture with four modules/ clusters (Visualization is done with “BrainNet Viewer” Xia et al. (2013) with empirical BOLD signals).

We further extend our analysis to understand the model’s performance in predicting the community structure for each participants. For this, an “average” functional connectivity has been developed, and the community structure of the brain network is evaluated after applying the Louvain algorithm for multiple iterations calculating the probability of two nodes residing in same community (Lancichinetti and Fortunato, 2012; Jeub et al., 2018). A similar methodology has been adopted for both simulated and empirical EEG and BOLD signals. Then the correlation coefficient between the two matrices is calculated (see Fig. 5(i)). In our earlier studies, we demonstrated that an oscillator-based model could retrieve information about the modular nature of the brain network embedded in the network topology and segregate the DMN from the rest of the network (Bandyopadhyay et al., 2023). Note that, earlier works (e.g. Castlado et al, 2023) also investigated the resting state network patterns based on K-means clustering, and not modularity (Castaldo et al., 2023). In the current study, we extend the model’s capacity to determine the static modularity, dynamic modularity, and representative community structure of the network from module allegiance matrix, which is derived from the empirical simultaneous EEG and fMRI data.

### Impact of Structural pruning on FC, FCD and community structure

A well-known method to understand structure-function relationship in a whole-brain model is to systematically perturb the connections in the structural connectivity (SC) and examine its effects in terms of functional connectivity (Cabral et al., 2012; Váša et al., 2015; Adhikari et al., 2015; Surampudi et al., 2018). A disruption in structure-function relationship has been shown to a key feature in neurological disorders, like-Schizophrenia (Stephan et al., 2009; Cocchi et al., 2014). In this in-silico perturbation technique, the structural connections between different ROIs are pruned systematically, either by proportional threshold or absolute threshold. The proportional threshold demonstrates the lesion of structural connectivity, retaining only a certain percentile of the strongest connections, and the percentile value serves as a threshold parameter, referring to the higher value corresponding to a more significant number of stronger connections present in the structural network. The perturbational impact of structural disconnectivity on the functional connectivity map is inspected both on static functional connectivity and dynamic functional connectivity. It is found that the correlation coefficient between simulated and empirical FC reduces as we increase the pruning of SC. A smaller proportion of stronger connections remains intact in the SC matrix with the decrease of threshold parameter in the proportional threshold, as illustrated in Fig. 6(b) for all the key band patterns of EEG data. Similarly, the *KS_distance_*, referring to the distance between the distribution of FCD values, increases with the increase of the structural lesion, as indicated in Fig. 6(a). This outcome reveals that, similar to FC, the structural lesion impacts the FCD or variability in FC, as determined from windowed EEG data for all the band patterns. Figures 6(e), 6(f), and 6(g) represent the histogram of FCD values depicting the impact of pruning on the functional activity of the brain. Even in clinical studies (Chu et al. 2015), it has been observed that FC evaluated from the source reconstructed EEG data is constrained by the SC estimated from probabilistic tractography, and the aberration in terms of pruning of SC significantly reduces the FC for all frequency bands (Chu et al., 2015). A similar outcome has been observed in the case of haemodynamic data —the virtual structural lesion has an inverse relationship with the predictive power of the model and a linear relationship with the *KS_distance_*, as demonstrated in Fig. 6(c) and 6(d), respectively. In our earlier work, a similar study was done to capture the impact of structural loss on functional connectivity maps in terms of FC and FCD of hemodynamic data (Bandyopadhyay et al., 2023). Interestingly, the current model can examine the impact of structural loss simultaneously on EEG and fMRI functional maps (both FC and FCD). In addition, the current study also investigates the “representative” community structure derived from the iterative community detection algorithm for each of the pruning conditions, i.e. both absolute and proportional thresholds. Figure 6(h) illustrates that the community structure obtained from the model, which incorporates pruned structural connectivity, shows a decreasing correlation coefficient between the community structures derived from simulated and empirical data as the proportional threshold is reduced. (Only cases of proportional thresholding are presented here; the cases of absolute threshold are discussed in section S2.2 of the Supplementary Document). It reflects that structural disconnectivity makes the model susceptible to losing its ability to capture the whole brain’s community structure. For both EEG and fMRI data, the impact of structural pruning on community structure emerged from simulation with pruned SC is shown in Fig. 6(h). It shows that with decreasing proportional threshold parameter, the SC network losses it’s ability to capture the community structure from the model. This phenomenon is shown by comparing the correlation coefficient between community structure network obtained from the simulated FC from the model with pruned SC in different cases, and the community structure obtained from the empirical data, and it is done for both electro-physiological and hemodynamic data.

**Fig. 6:**
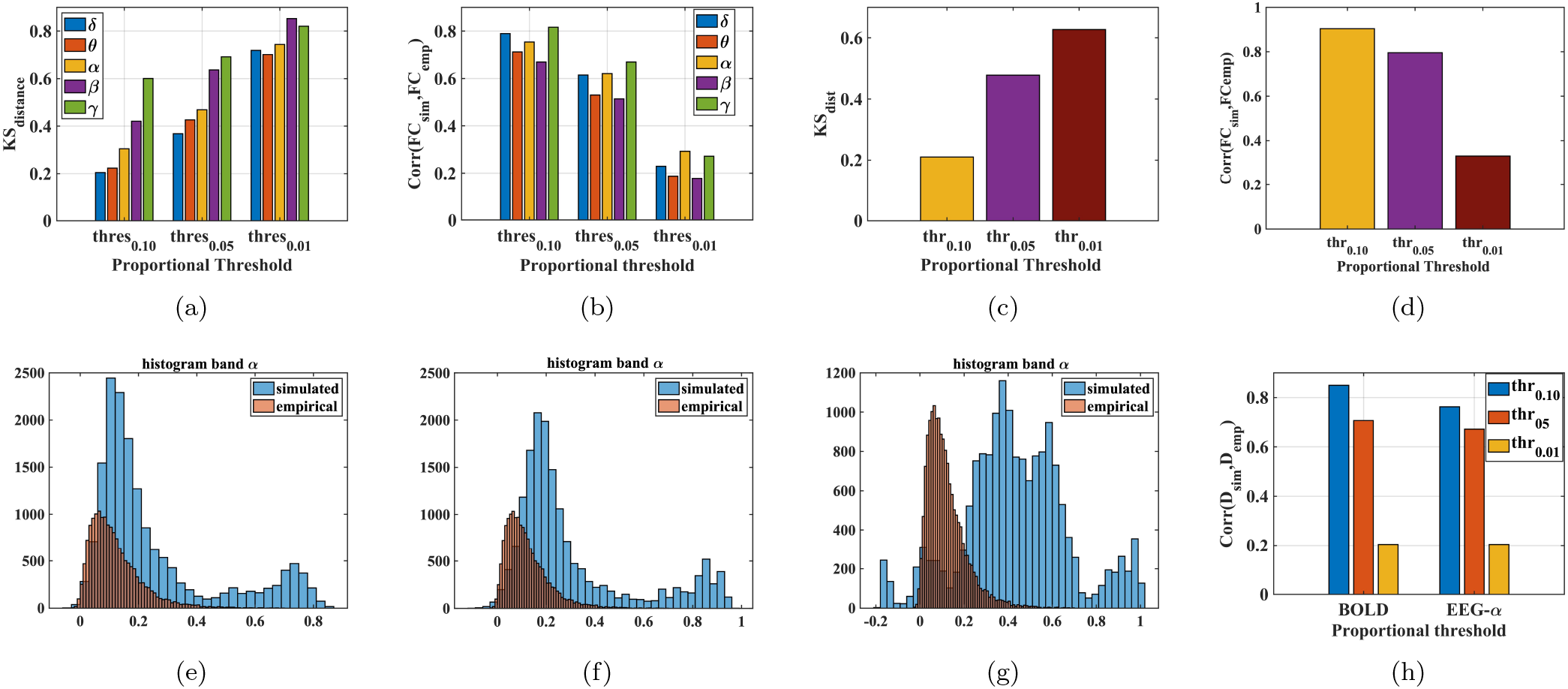
Inferring the effect of structural pruning in the mathematical framework for EEG-fMRI concurrent data simulation in terms of FC, FCD, and community structure. Fig. (a) shows the effect of pruning by proportional threshold in FCD calculation for all the frequency bands of EEG. Fig. (b) compares the effect of pruning by proportional threshold in calculating the FC for all the frequency bands of EEG. Figs. (c) and (d) demonstrate similar outcomes for trimming the structural connectivity matrix. Figs. (e), (f) and (g) refer to the histogram of the FCD values, indicating the difference in the distribution of the FCD values for simulated and empirical data for EEG-*α*. Fig. (h) refers to the difference in community structure for different pruning cases, where the structural pruning is done with decreasing order of threshold parameter. It is shown for both the cases of EEG (EEG-*α*) and BOLD signals.

### Perturbational spatio-temporal Integration

The small-world architecture of the human brain, with its modular yet integrated structure, manifests a unique balance in segregation and integration, rendering seamless information processing, providing characteristic cognitive behaviour and different consciousness states (Luppi et al., 2019, 2021). Previous findings demonstrated that structural disconnection disrupts the brain’s integration and segregation state, with induced randomness causing a reduction of integration level and formation of isolated clusters that enhance segregation (Deco et al., 2015; Lord et al., 2017). However, such information has been gleaned from analyzing the topological behaviour of the human brain, either by looking at the modularity value or evaluating the largest connected component of the graph (Cabral et al., 2012; López-González et al., 2021). Integration and segregation in the spatio-temporal domain have not been studied well, and the current model has the ability to predict the integration and disintegration in macro-level due to structural lesions. At this end, we have investigated the impact of structural loss both in different temporal scale data with different frequency ranges, i.e. different frequency band patterns of electro-physiological data, and the haemodynamic data, providing the unique opportunity of conducting in-vivo perturbation and corroborate with the experimental data from simultaneous extraction of EEG and fMRI signals. A more detailed discussion about the integration and segregation state is given in the “Methods” section. At this end, the spatiotemporal integration for BOLD data is achieved by evaluating the cartographic profile at each sliding window, and these profiles have been clustered into two groups according to participation coefficient value-predominantly integrated and predominantly segregated (Shine et al., 2016). The duration of the predominantly integrated state as a proportion of the total time points is referred to as the Integrated State of Occurrence Rate (ISOR) (Wei et al., 2022). Notably, the evaluation of the integration state from EEG data for different frequency bands is based on spectral graph theory, particularly the nested spectral partition (NSP) method (Zhou et al., 2023; Wang et al., 2022). Figure 7(b) demonstrates the ISOR for simulated, empirical and the different structural lesion cases. It is shown that the ISOR is decreased monotonically with the decrease in the proportional threshold. A similar result can be obtained with the absolute threshold of the SC (given in the section S2.2.4 in the supplementary document online). Interestingly, in the current work, we restricted in-silico perturbation to random structural pruning, not a selected or focal lesion. However, it has been shown that such a lesion can unravel the importance of individual node’s topological features in global integration and small-world networks (Wei et al., 2022). It has already been shown that ISOR can be a perfect repertoire of cognitive disorders and behaviour (Shine et al., 2016; Luppi et al., 2019). Figure 7 (c) reveals that the model can capture the integration level (*M_in_*) of the brain observed from the electro-physiological data for different frequency bands, and interestingly, it is shown that the structural loss causes a reduction in integration level across all frequency bands in EEG monotonically. This simulated outcome can be supported by the recent imaging data and signal analysis by Handiru et al. (2021) on chronic TBI participants, where it was found that the structural aberration and damage cause the increase in segregation and reduction in integration (Shenoy Handiru et al., 2021).

**Fig. 7:**
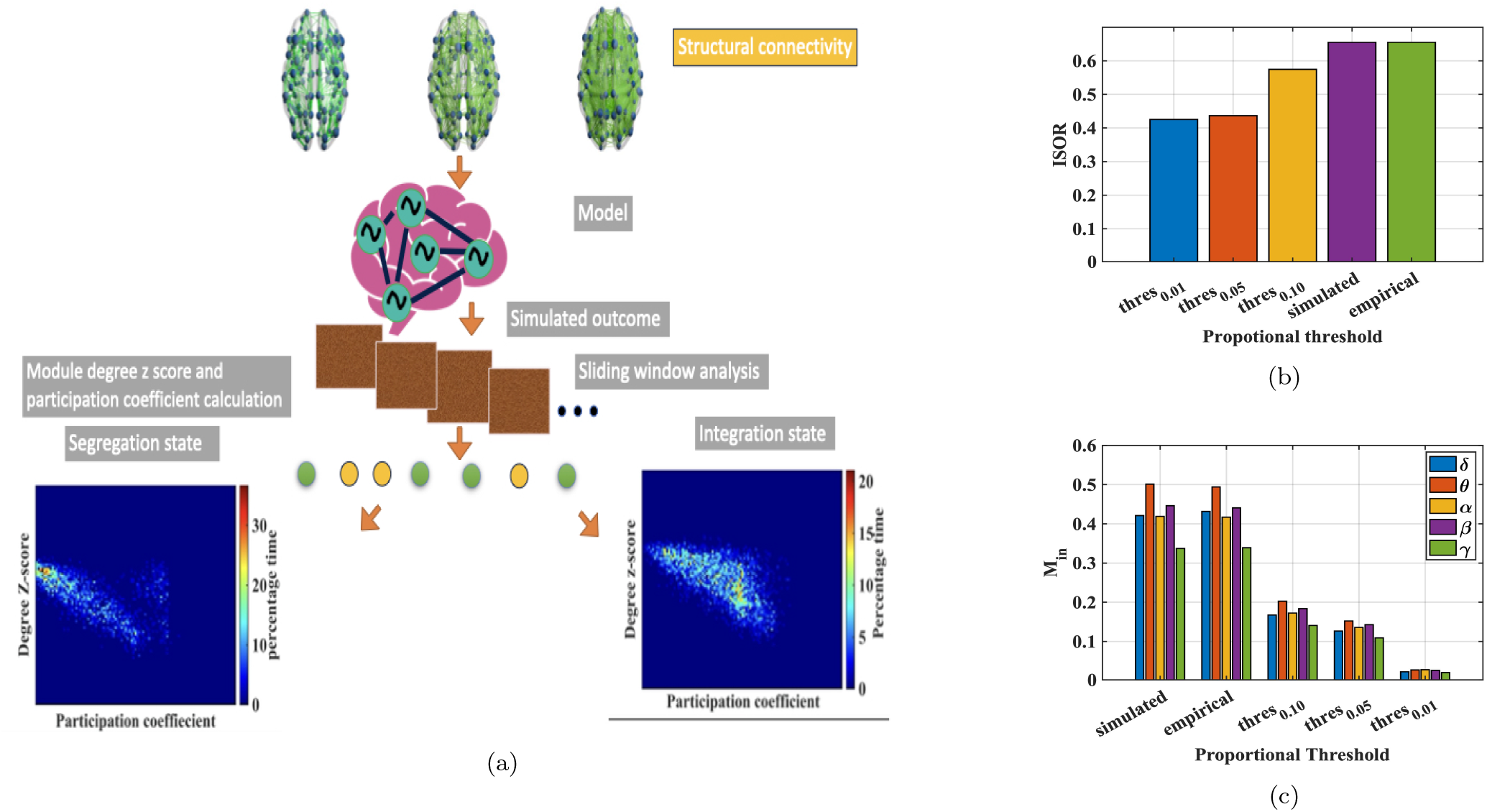
Spatio-temopral Integration and segregation. Fig. (a) illustrates a schematic diagram of evaluating the integration state by taking the cartographic profile for each time window. Fig. (b) and (c) shows the impact of structural pruning by proportional threshold on integration state of whole brain for heamodynamic and electro-physiological data, respectively.

### Application of the model to alternative dataset

The current model has also been applied to another dataset, which includes structural data and simultaneous EEG-fMRI data for the sake of validation (Deligianni et al., 2016)[Link to the database-https://osf.io/94c5t/]. The pre-processing steps have been discussed in detail in the supplementary section. The repetition time of the fMRI signal is 2.16 seconds and the sampling frequency of the EEG signal is 250 Hz. An approach similar to the one described in the “Basic Mathematical Model” is adopted to simulate this validation dataset. At this end, we will describe how this model can capture the FC, FCD of the validation dataset. The results described here are for first indexed data in the “EEG-fMRI-NODDI” dataset. The Fig. 8(e) shows the ability of the current framework to mimic the FC. All the FC for different frequency bands along with FC from BOLD signals can be captured with this model. Similarly, the model can capture the FCD also which can be evaluated by the distribution of FCD matrix values, which is evident from the Figs. 8(f). In Figs. 8(g) and 8(h), the histogram of FCD values for both simulated and empirical data demonstrates that the concurrent EEG-fMRI model can reproduce the FCD values obtained from empirical data. Note that, Figs. 8(a) and 8(b) represent the FCs deduced from simulated and empirical data for BOLD signals, respectively. Furthermore, the AEC FCs for alpha frequency bands for simulated and empirical are given in Figs. 8(c) and 8(d).

**Fig. 8:**
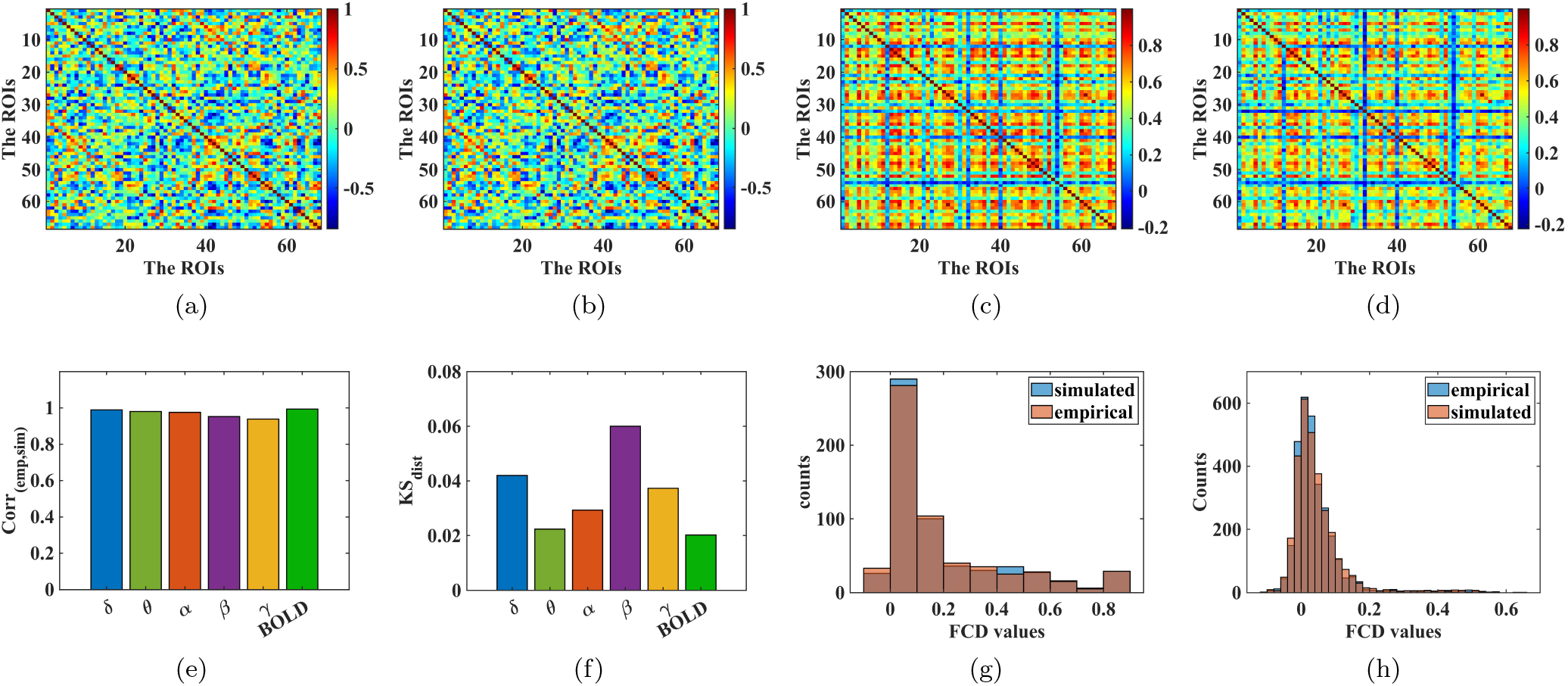
Simulated outcome for validation dataset. Figs. (a) and (b) individually represent the simulated and empirical BOLD signals’ functional connectivity matrix. Figs. (c) and (d) reveal the AEC based FC for simulated and empirical EEG data of alpha frequency band region; Fig. (e) presents the predictive power of the model for the first participant of the EEG-fMRI-NODDI dataset. Fig. (f) reveals the Kolmogorov-Smirnov (*KS_dist_*) distance between the distribution of the FCD values for simulated and empirical BOLD signals, and EEG signals for different frequency bands. Figs. (g) and (h) represent the distribution of the dynamic functional connectivity values for both the simulated and empirical data of *EEG* − *α* and BOLD

## Discussion

Recent advances in neuroimaging tools and techniques have created an unprecedented opportunity to understand neural activity in the brain at multiple temporal scales. Simultaneous recording of EEG-fMRI data poses the challenge of understanding the brain at two different frequency bands simultaneously – the lower frequencies of fMRI (around 0.01 Hz) and the faster frequencies of EEG (0.1Hz − 50 Hz in this study). Simultaneous EEG-fMRI recording has emerged as a valuable tool for deciphering the neural basis of the hemodynamic response and providing insights into behavioral data (Scrivener, 2021; Warbrick, 2022). However, extracting high-quality data can be challenging due to artifacts induced during simultaneous recordings, particularly MR-induced artefacts (Bullock et al., 2021). Recent developments in MRI-informed source localization enable the mitigation of the inherent volume conduction/spatial source localization problem, leading to the possibility of decoding the underlying relationship between the fast-spiking temporal behavior of EEG and its delayed hemodynamic response at longer time scales (Warbrick, 2022; Jorge et al., 2014). Multimodal recording has important clinical applications, including localizing epileptogenic zones (Sadjadi et al., 2021), tracking the propagation of interictal epileptiform discharge (IED) (Grouiller et al., 2011; Vulliemoz et al., 2011), and even identifying the cognitive malfunction associated with it (Shamshiri et al., 2019). The spatiotemporal patterns generated from concurrent EEG-fMRI data can be useful for multiple fields, such as neuro-feedback and focal brain stimulation (Mano et al., 2017; Warbrick, 2022). MRI-assisted precise localization of neural activity data, such as EEG data, combined with structural connectivity (SC) information and fMRI data, has augmented the development of computational models of concurrent EEG-fMRI that help to understand the underlying relationship between two different neuroimaging modalities. (Schirner and Ritter, 2023; Breakspear, 2017). Over the past decade, various mathematical models have been developed, ranging from biologically grounded models to those based on machine learning and abstract conceptual approaches. The proposed modeling approach stands midway between detailed biophysical models of brain dynamics and more abstract models. To this end, we compare the proposed model with other existing models based on modeling features and the accuracy of prediction. Some of the most popular models in this domain are given in Table 1, along with their key features, accuracy, and clinical significance. Note that the described models are simulated on different datasets. Readers are requested to refer to the supplementary document for more information about these models. The model presented by Schirner et al. Model 1 in Table 1 is more biologically plausible since it is based on an earlier mean-field model that incorporates the firing rates of excitatory and inhibitory neurons and the coupling between excitation and inhibition (Schirner et al., 2018). Several insights can be drawn from this model, such as the relationship between alpha wave and low-frequency BOLD signals and the importance of excitation-inhibition balance.

**Table 1.** Comparison of models integrating EEG and fMRI modalities.

| Model | Cross-Modality | Model Description | Dataset | Power | Clinical Importance |
| --- | --- | --- | --- | --- | --- |
| Model 1 (Schirner et al., 2018) | EEG-fMRI | Modified neural mass model | Berlin | 0.50 (approx) | Relationship between Alpha wave and neural firings. |
| Model 2 (Mann-Krzisnik and Mitsis, 2022) | EEG-fMRI | Hopf-oscillator-based network model with amplitude coupling | EEG-fMRI-NODDI, Motor | 0.28 (approx) | Effect of physiological confounds (e.g., cardiac rhythm) are investigated. |
| Model 3 (Castaldo et al., 2023) | fMRI-MEG | Wilson-Cowan and Hopf oscillator model | Own dataset | 0.56 (approx) | Features include synaptic plasticity, identification of RSNs, and metastable oscillatory modes |
| Model 4 (Ahmed et al., 2022) | EEG-fMRI | Reaction-diffusion system with Wilson-Cowan model | EEG-fMRI-NODDI | 0.70 (approx) | Structural disconnections and their effects are investigated. |
| Model 5 (Glomb et al., 2020) | EEG-fMRI | Empirical signals are processed (smoothed) using the SC graph. | Own dataset | 0.60 (approx) | Community structure of EEG and fMRI |
| Model 6 (ours) | EEG-fMRI | Discussed in the "Methods" | EEG-fMRI-NODDI, Berlin dataset | 0.97 (approx) | Community structure, impact of structural loss on FC, FCD, functional integration, relationship between EEG-fMRI |

Castaldo et al., in their recent study, used a Wilson-Cowan-based mass model to explore the impact of conduction delay and homoeostatic plasticity, demonstrating the balance between excitation and inhibition mechanisms in whole brain dynamics (Castaldo et al., 2023). It is important to note that in model 3, as referred to in Table 1, the BOLD signal was generated using the Balloon-Windkessel model, which produces haemodynamic response based on the neural activity produced by the neural mass model. Meanwhile, Hopf oscillator based network model has been adopted to capture MEG data, where each oscillator has negative-valued bifurcation parameter and 40 Hz intrinsic frequency. Conversely, the model identified as Model 5 in Table 1 proposed by Glomb et al. (2020), emphasizes the significance of structural connectivity, and suggests that the functioning of an individual node or region of interest (ROI) is primarily governed by that specific node and the collective activity of its neighboring nodes that are directly connected by white matter tracts. (Glomb et al., 2020). This work illustrates the relationship between EEG and BOLD signals at the macroscale level, considering the community structure and the correlation between EEG-FC and BOLD-FC. However, most of the above discussed models do not capture the concurrent EEG-fMRI data in a single modelling framework, making it challenging to determine the relationship between the slow hemodynamic signal and the fast electrophysiological signal. Furthermore, the models exhibit lower predictive power compared to the current model.

Apart from the modeling approach of Krzisnik and Mitsis (2023), referred to as Model 2 in Table 1, all the other models in the Table 1 simulated the electrophysiological and hemodynamic signals separately. Therefore, the model developed by Krzisnik and Mitsis comes closest to our modeling approach (Mann-Krzisnik and Mitsis, 2022). The model is based on extracting the dominant components with canonical polyadic decomposition (CPD) of the Hemodynamic response function, which are evaluated in the spatial and spectral domains as a response function estimated from LFP/EEG data and BOLD signals. For their modeling approach, two sets of oscillators, LFOs, and HFOs, are taken with two different frequency components, i.e., LFOs with fixed intrinsic frequency at 0.08 Hz and HFOs are set at 2 Hz and 10 Hz for the left and right hemispheres individually. There are striking differences between such a model and the model developed in the current study. The current model has a frequency learning algorithm so that the oscillators iteratively learn the spectral components of the corresponding teaching signal. Learning the amplitude of the complex nature of power coupling by the Hebbian learning algorithm also provides a chance to underscore the connectivity pattern between LFOs and HFOs, which resembles the relationship between EEG and fMRI. Apart from that, the current model has outperformed all the other models regarding predictive power.

This is the first model that captures the simultaneous impact of structural dysconnectivity at the functional level at two distinct temporal features obtained by EEG and fMRI. Such a study renders a comprehensive understanding of the importance of structural connection and its lesion in the functional connectivity map, community structure, and integration and segregation behavior of the brain at the spatiotemporal scale. Many experimental and theoretical studies have been done to inspect the impact of structural lesions on electrophysiological data and hemodynamic data individually; however, the current framework can integrate these experimental procedures within a mathematical framework (Babaeeghazvini et al., 2021). Another important application of such a model would be on possible rehabilitation strategies for neurological disorders, i.e., with the help of deep brain stimulation (DBS) and trans-cranial direct current stimulus (tDCS). Already, networks of Hopf-oscillators have shown promising outcomes in understanding the impact of non-invasive brain stimulation, which can be pivotal for planning experimental protocols, i.e., montage selection, amount of stimulation required, etc (Bandyopadhyay et al., 2023; Saenger et al., 2017; Deco et al., 2019). However, few clinical studies have been done on simultaneous EEG-fMRI combined with non-invasive brain stimulation, such as transcranial alternating current stimulation (tACS). Or transcranial magnetic stimulation (TMS). They yield valuable insights into brain dynamics and have potential applications in rehabilitation. (Clancy et al., 2022; Peters et al., 2020).

Cabral et al. have delineated three benchmark parameters to validate any whole brain model: spatial, temporal, and spectral (Cabral et al., 2017). Spatial validation refers to the predictive power based on FC calculation and capturing the resting state network. The model’s high prediction power and evaluation of modularity and corresponding community structure in spatial and spatiotemporal domains satisfy this criterion. FCD analysis fulfills the temporal validation and compares the simulated EEG and BOLD signal data with the empirical one. Typically, for electrophysiological data, time-windowed amplitude envelope correlation (AEC), and for hemodynamic data, Pearson’s correlation-based functional connectivity (FC) is employed with a sliding window technique. The power spectral data of individual ROI is compared with the corresponding empirical data for spectral validation (see Figs. 2(d), and 2(i)). However, Bassett and colleagues revised these criteria into three broader categories: descriptive, explanatory, and predictive validity (Bassett et al., 2018). Descriptive validity includes approximating simultaneous EEG-fMRI data using FC, FCD, and modularity-based validation measures, and the model successfully fulfills it. Furthermore, the model is employed to decipher the empirical relationship between FC fMRI and FC derived from different frequency bands of EEG signals from a pure modeling perspective, leveraging the Hebbian learning rule deployed to the power coupling between HFO and LFO oscillators, which approves that the model can explain the empirical phenomena. To validate the prediction, an in-silico perturbation technique characterized by randomly pruning the SC matric has been adopted in this study, and its effect on FC, FCD, and integration measurement is divulged here. To showcase the superiority of the current model, a few of the models described in the Table 1 are simulated on the Berlin data (2018) (Schirner et al., 2018), and the EEG-fMRI-NODDI dataset given by Deligianni et al. (2016) (Deligianni et al., 2016). Noteworthy, some of the models have been built earlier for demonstrating either electrophysiological or hemodynamic data. Fig. 9(a) shows a schematic figure highlighting the differences among various modeling approaches - Hopf oscillator model with delay and fixed frequency (Castaldo et al., 2023) (Castaldo et al., 2023), Hopf oscillator model without delay and inhomogeneous frequencies (Deco et al, 2017) (Deco et al., 2017b), and neural mass model representing excitation and inhibitory firing of neurons (Castaldo et (2023), Abeysuriya et al.(2017)) (Castaldo et al., 2023; Abeysuriya et al., 2018). In Fig. 9(b) shows the performances of the models provided in the Table 1, and the models are performed on the two aforementioned datasets (“EEG-fMRI-NODDI” and “Berlin dataset”). Some of the models have already been applied to these data sets, and performances were reported. For comparing the models for capturing the hemodynamic data, the model given by Deco et al. (2017) is performed on the homogeneous setting, referring to oscillators’ amplitude being kept constant with changing global connectivity. The highest prediction power with optimized parameter setting is found to be 0.35. Similarly, Model 3 (as per Table 1) with optimized parameter setting estimates the prediction power at 0.38 for the Berlin dataset to model solely the fMRI data. Subsequently, to model the EEG data, a network of delayed Hopf oscillators is applied; it provides the predictive power of around 0.39 for all the channels, which is quite comparable to the earlier reported studies. On the other hand, the dataset provided by Daligieni et al. was already simulated on Model 2 and reported the prediction power at 0.30 (ranging from 0.24 − 0.32). Model 4, a reaction-diffusion system-based model, achieves a prediction power of 0.71 (approx.; varies for different band patterns of EEG and fMRI) when applied to the EEG-fMRI-NODDI dataset. Similarly, for EEG modeling, Abeysuriya et al.’s neural mass model is applied to the Berlin dataset with optimized parameters (parameters are given in the Supplementary document).

**Fig. 9:**
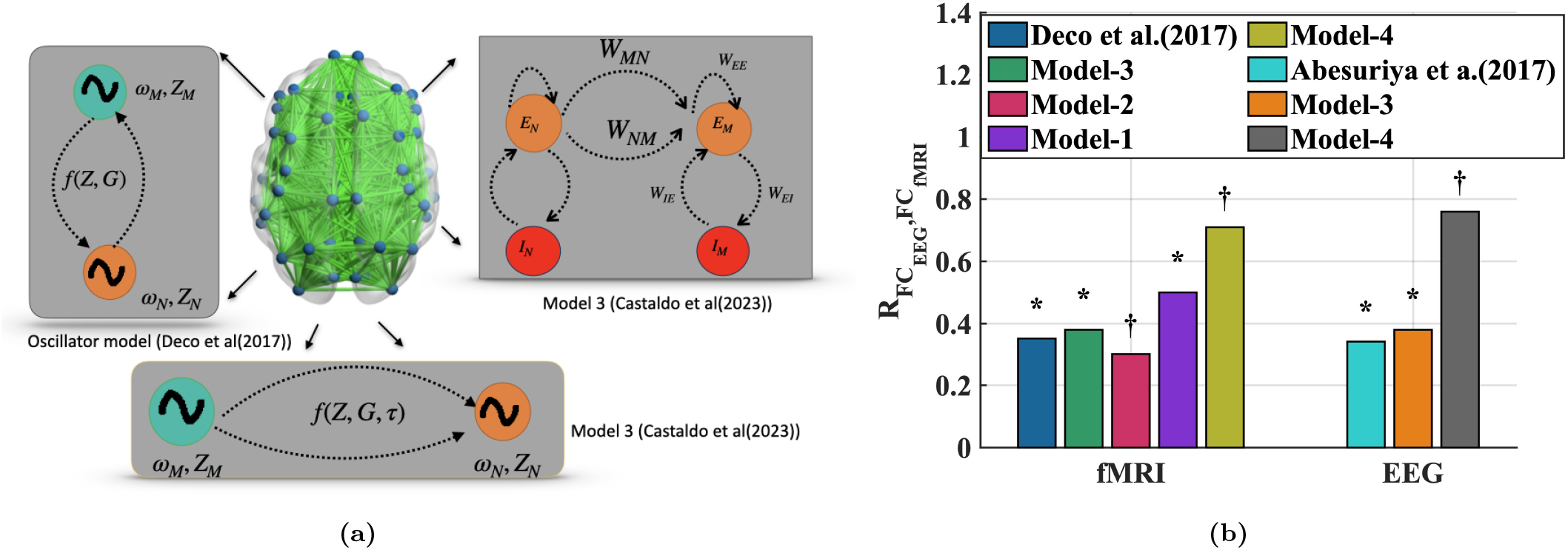
Comparison of models integrating EEG and fMRI modalities, focusing on predictive power and clinical importance. Figure (a) provides a schematic diagram for three different models, including neural mass models, networks of real-valued coupled oscillators, and oscillators with real-valued coupling and time-delay (*τ*). Figure (b) illustrates the predictive power of different models and approaches.* and † represent Berlin and EEG-fMRI-NODDI datasets, respectively.

It is important to note that one of the major limitations of the current model is that it is purely phenomenological, disregarding the phenomenon of cerebral blood flow constrained by the neural activity orchestrated by the energy-driven cascades of biological events in the neuron-astrocyte-blood vessel at a microscopic level, which is typically referred as neurovascular coupling (Iadecola, 2017). However, the underlying biological and metabolic events are not fully understood; multi-compartment forward models have showcased promising outcomes in illustrating the biological events behind the BOLD signal (Mathias et al., 2018). Nonetheless, such models are devoid of inferring the brain at the macroscopic scale. For future directions, the mechanistic model discussed in the current study can be extended to incorporate the information on neural dynamics-driven vascular activity at the population level, which can provide the opportunity to understand the neurovascular coupling event on a macroscopic scale.

Multilayer networks of oscillators have been in the literature domain in modelling to understand different b iological phenomena, cognitive behaviour, and even in visual information processing (Wang and Terman, 1997; Borisyuk et al., 2002; Li, 1998). Such a multi-layered oscillator modelling framework, where an unsupervised learning algorithm teaches the connectivity pattern between two different s ets o f o scillators, c an b e e asily e xtended t o u nderstand t he u nderlying r elationship b etween t he d ata e xtracted f rom multi-modal instruments, which is not limited to the EEG-fMRI, and can be extended to other applications, like-the brain-gut model from EGG (Electro-gastriogram)-fMRI, the brain-heart model with ECG(Electrocardiogram)-fMRI etc. One of the major drawbacks of the current modelling approach is to understand the brain as a separate entity from the body by discarding the visceral signal emerging from different organs. An integrated, wholesome approach taking leverage from the multilayer networks of oscillators can illustrate brain-body interaction.

## Data Availibility

Two sets of dataset are used in this modeling study. The “Berlin dataset” is available with necessary permission and documentation from the online repository-https://osf.io/mndt8/; the “EEG-fMRI-NODDI” dataset is available at following online repository-https://osf.io/94c5t/.

## Code Availability

The codes (MATLAB scripts) used for simulation and producing the results are available upon reasonable request from the corresponding author.

## Competing interests

No competing interest is declared.

## Acknowledgment

we like to thank the authorities of Department of Biotechnology, IIT Madras for their constant support. We like to acknowledge professor Dr. Petra Ritter from Charite University Medicine Berlin for granting permission to avail the “Berlin” dataset.

## Author contributions statement

A.B. conducted all the simulations and wrote the original draft; A.B., V.S.C and D.R. analysed the results. The work is conceptualized and planned by V.S.C. who edited and reviewed the manuscript. V.S.C, and D.R provided valuable suggestions to update the results and improve the manuscript. All authors have contributed in writing the manuscript and reviewed and finalized the manuscript.

### Supplementary Documents

Supplementary data is included in the manuscript pdf.

## Funding

This research work received no external funding.

## Supplementary Document

### S1 Detailed Methodology

#### S1.1 Mathematical framework

The equations that describe training and activation dynamics of the oscillator network used in this paper were described earlier in relation to various applications, i.e. modeling tonotopic maps [Biswas et al., 2022], approximating EEG signals for a limited number of channels [Biswas et al., 2021], understanding the relation between brain structure to function with the help of BOLD signals from Functional Magnetic Resonance Imaging (fMRI) and tractography data from Diffusion Tensor Imaging(DTI) study [Bandyopadhyay et al., 2022]. Here we will briefly describe the governing equations for first stage and second stage of learning.

##### S1.1.1 First Phase of Learning

As discussed in the main text, the current model is composed of two layers of the oscillatory network (LFO and HFO) corresponding to the fast and slow oscillations of EEG and fMRI, respectively. We noticed that such a model’s performance is strongly dependent on the number of oscillators in each cluster or sub-cluster (*n_HF O_* or *n_LF O_*). The first stage is a linear network, where the oscillators of each ROI connect to form a network according to its physical connectivity as assigned by the structural connectivity (SC) network. In the first stage of learning, we learn the oscillators’ intrinsic frequencies (*ω_LF O_*, and *ω_HF O_*) and the corresponding phase differences using a modified form of Hebbian learning (eqn. 3). Later, the inter-layer (between *LFO* and *HFO*) connections are trained with this modified Hebbian learning.

The corresponding equations are as follows-

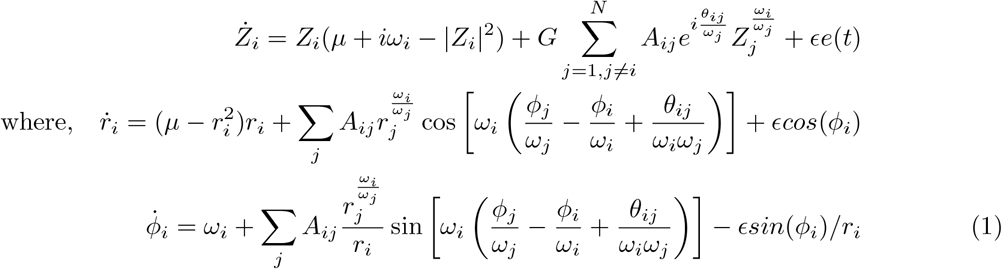

The forward connections and frequency learning are given by - [Biswas et al., 2021,Bandyopadhyay et al., 2023],

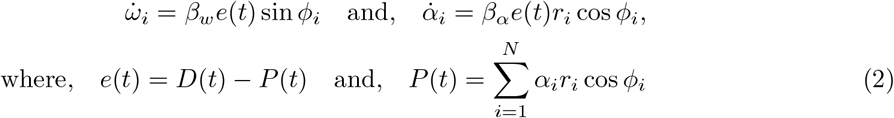

and, for Hebbian learning, the modified form looks like –

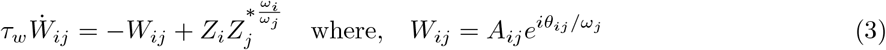

For updating the amplitude of the Hebbian learning, the amplitude, |*W_i,j_*| is kept in a bounded region defined by |*W_i,j_*| ∈ [0.005, 0.05] for LFO-HFO connections. Note that, in the main text, we define the connections between LFO and HFO oscillators as *W_if,je_*. There are two other nature of connections are present, i.e. connections in-between the sub-cluster, or |*W_ir,jr_*| and the connections between different sub-clusters of LFOs or HFOs (inter-cluster connections) representing two separate ROIs, i.e. |*W_ir,js_*| as given in the main text. The values assigned to them are provided in Table S1.

##### S1.1.2 Second phase of Learning

The second phase of learning pertains to learning the EEG signals with a complex-valued feedforward network, with a hidden layer of twenty sigmoid neurons, are used to train the network. Previous findings also suggest that such network is dependent on number of epochs and the neurons in the hidden-layer [Bandyopadhyay et al., 2022]. For details of the learning architecture, readers may also refer to the main text and the Supplementary section of a previous study [Bandyopadhyay et al.,2023].

The loss function utilized here is a squared error loss, given by [Bandyopadhyay et al., 2023]-

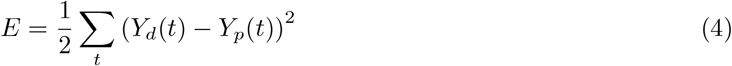

Here, *Y*_*d*(*t*)_ and *Y*_*p*(*t*)_ are the simulated and empirical signals (for both EEG and fMRI) for each ROI. There are an ensemble of oscillators corresponding to each ROI; the oscillators are connected by SC matrix i.e only oscillators pair with other oscillators corresponding non-zero SC weight are connected. In the feedforward network whose output neurons model the empirical signals, there is a hidden layer of sigmoid neurons between the oscillator corresponding to the ROI and the output neuron (see Fig. 1.(c) in main text).

The oscillator-layer, with a dimension of *N* ×*T*, is represented by *Zhbs* ∈ ℂ^*N* ×*T*^, where N denotes the number of oscillators, and T is the number of simulated time points. The weights (*W* 1*bs* ∈ ℂ) are also complex in nature, which are updated by the back-propagation. The forward connections between the oscillators and the hidden layer are given by –

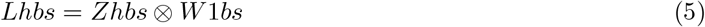

The activation function is given by –

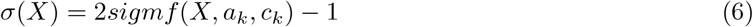

where, *sigmf* is sigmoidal membership function, represented by-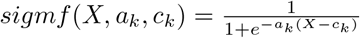, where, *a_k_* = 0.5 and, *c_k_* = 0

The output of the hidden layer is given by —

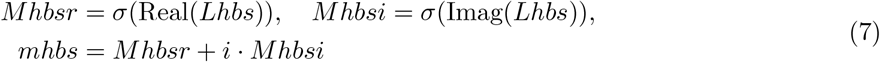

The transformation from the hidden layer to the output layer is given by —

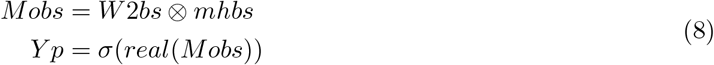

Note that, only the real part is taken for further analysis. *W* 2*bs* is a complex valued weight(*W* 2*bs* ∈ ℂ). The backpropagation rules are already given in the Supplementary sections in the previous work [Bandyopadhyay et al., 2023]. The learning rate of the update of the weights are fixed at the rate of 0.001 (see the Table S1).

#### S1.2 Parameter Table

**Table S1:**
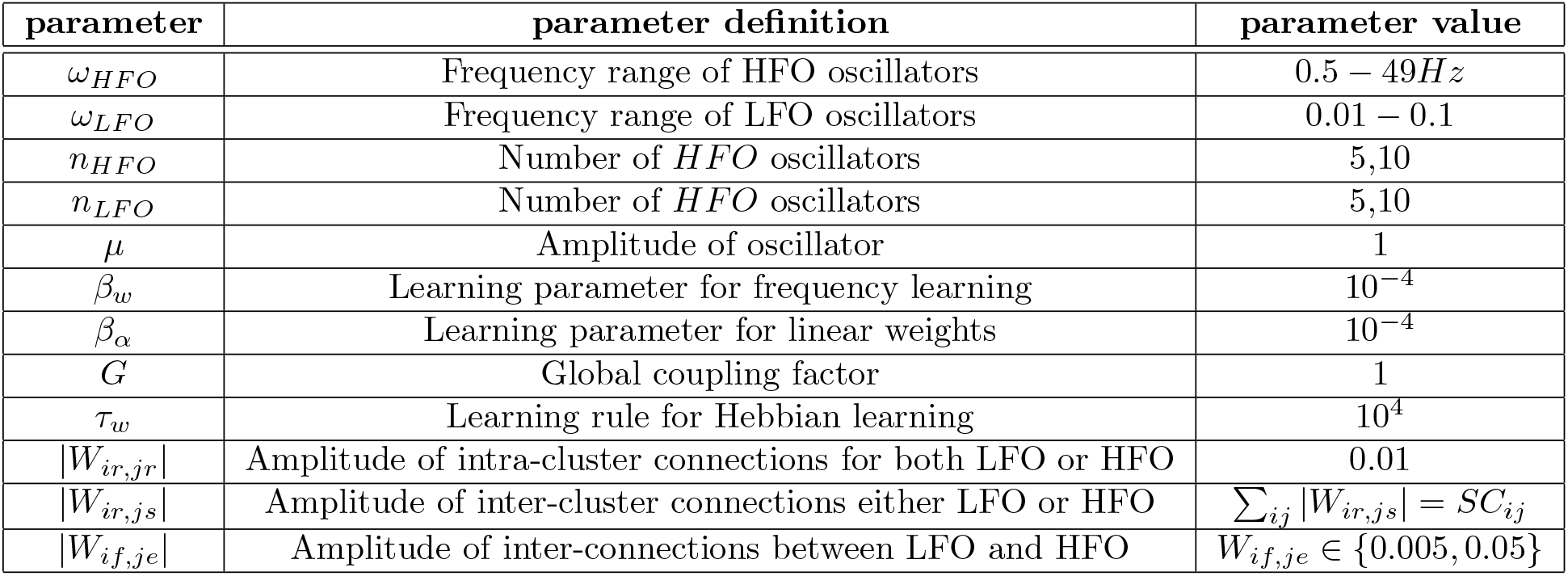
Parameter table depicting some of the parameter values taken in this study.

#### S1.3 Data Processing step of the Validation Dataset

Model validation is done by the ‘EEG, fMRI and NODDI dataset’ [Deligianni et al., 2016]. This dataset is popular in modeling domain, particularly for analyzing simultaneous EEG-fMRI data. We now provide a brief description of the acquisition of the data, pre-processing steps, and the required imaging toolboxes to analyze the raw data. The dataset was acquired from seventeen adult participants with including 11 males and 6 females from the age group of mean 32 years and standard deviation 8 years. The scanning parameters are as follows-TR-2160 ms, TE-30 ms, effective voxel size-3.3 × 3.3 × 4.0 mm, and flip angle-75^°^. For more about the MRI set up, the readers may refer to the dataset repository and the https://osf.io/94c5t/ related article [Repository can be found here-[Deligianni et al., 2016].

For structural data pre-processing, FREESURFER toolbox were utilized where the anatomical data (T1-weighted image) were processed with the default preprocessed stages (pipelines), which were already described in detailed in earlier works and the documentation of Freesurfer [Fischl, 2012]. These procedures have been found to be more accurate to segmentalize the grey matter and white matter’s boundary. Cortical grey matter has been further parcellated based on Desikan-Killiany atlas as given in the Table S2.

For, fMRI data processing, the CONN toolbox was used [Whitfield-Gabrieli and Nieto-Castanon, 2012]. However, in this implementation, instead of volumetric analysis, the analysis was done on surface space where the preprocessed structural data obtained from the FREESURFER toolbox, as discussed earlier, were utilized. We followed this pre-processing pipeline as outlined in Andy’s Brain Book (Jahn, 2022. doi:10.5281/zenodo.5879293). The processed data were also filtered with a band-pass filter of 0.008 − 0.08 Hz as default setting in CONN toolbox.

Structural connectivity data is already given in the “EEG, fMRI and NODDI dataset” repository. Some of the important scanning parameters are - TR-8300 ms, TE-98 ms, flip angle-15^°^, effective voxel size - 2.5× 2.5 × 2.5 mm etc. For other parameters readers may refer to the database repository and the original article [Deligianni et al., 2016].

EEG data were acquired with a 64-channel embedded electrode cap, which was MR compatible. The EEG data was already preprocessed with the help of Brain Vision Analyzer 2 (BrainVision Analyzer, Version 2.2.2, Brain Products GmbH, Gilching, Germany). Later, source reconstruction in MRI’s structural space defined by the Desikan-Killiany Atlas is done by the Brainstorm toolbox [Tadel et al., 2011]. The pipeline for this procedure is detailed in the Brainstorm documentation and was followed step-by-step in this study. A similar procedure was employed in earlier works to extract source reconstructed EEG signals [Schirner et al., 2018]. Realistic forward head models (along with scalp and skull) are constructed using the boundary element model by OPENMEEG [Kybic et al., 2005]. For inverse solution, standardized low-resolution brain electromagnetic tomography, or sLORETA in short, was used to extract the source activity distributed in the cortical surface, which is generated by the aforementioned BEM head model where a cortical surface is created by triangulated mesh, and each vertex resembles a perpendicular dipole generator representing neural activity [Pascual-Marqui et al., 2002].

#### S1.4 Desikan-Killiany Atlas

**Table S2:** Brain Regions in Left and Right Hemispheres.

| Index 1-34, Left | Index 35-68, right |
| --- | --- |
| 1. bankssts | 35. bankssts |
| 2. caudalanteriorcingulate | 36. caudalanteriorcingulate |
| 3. caudalmiddlefrontal | 37. caudalmiddlefrontal |
| 4. cuneus | 38. cuneus |
| 5. entorhinal | 39. entorhinal |
| 6. fusiform | 40. fusiform |
| 7. inferiorparietal | 41. inferiorparietal |
| 8. inferiortemporal | 42. inferiortemporal |
| 9. isthmuscingulate | 43. isthmuscingulate |
| 10. lateraloccipital | 44. lateraloccipital |
| 11. lateralorbitofrontal | 45. lateralorbitofrontal |
| 12. lingual | 46. lingual |
| 13. medialorbitofrontal | 47. medialorbitofrontal |
| 14. middletemporal | 48. middletemporal |
| 15. parahippocampal | 49. parahippocampal |
| 16. paracentral | 50. paracentral |
| 17. parsopercularis | 51. parsopercularis |
| 18. parsorbitalis | 52. parsorbitalis |
| 19. parstriangularis | 53. parstriangularis |
| 20. pericalcarine | 54. pericalcarine |
| 21. postcentral | 55. postcentral |
| 22. posteriorcingulate | 56. posteriorcingulate |
| 23. precentral | 57. precentral |
| 24. precuneus | 58. precuneus |
| 25. rostralanteriorcingulate | 59. rostralanteriorcingulate |
| 26. rostralmiddlefrontal | 60. rostralmiddlefrontal |
| 27. superiorfrontal | 61. superiorfrontal |
| 28. superiorparietal | 62. superiorparietal |
| 29. superiortemporal | 63. superiortemporal |
| 30. supramarginal | 64. supramarginal |
| 31. frontalpole | 65. frontalpole |
| 32. temporalpole | 66. temporalpole |
| 33. transversetemporal | 67. transversetemporal |
| 34. insula | 68. insula |

#### S1.5 Different comparison models

In the Discussion section of the main text, we compared the current model with the earlier developed models. However, as discussed earlier, such models were developed to model data from a single modality, such as empirical data from EEG, fMRI, or MEG. Nonetheless, it would be interesting to compare some of the above models, since the modelling approaches are quite similar,i.e, modelling the whole brain using a network of Hopf oscillators, constrained by structural connectivity. Also, a few recent works, like-Castaldo et al. [Castaldo et al., 2023], used different oscillator models (Hopf-oscillator based model and Wilson-Cowan model) to simulate concurrent MEG-fMRI data. In Table 1 (Fig. 9(c)), in the main text, we described the modelling strategy and it’s accuracy for a few recent modelling studies. However, they are often simulated and validated with their own dataset. So, it becomes imperative to test those models with “Berlin” and “EEG-fMRI-NODDI” dataset, which corroborate the current modelling scheme. Our aim is to show that the current model exhibits the superiority in terms of approach (without the rigorous parameter optimization technique) and modelling accuracy in terms of approximating empirical FC, FCD and modular architecture of brain. In our previous study, we highlighted the importance of power coupling between two or more Hopf oscillators in order to build a large-scale whole brain model with Hopf oscillators [Bandyopadhyay et al., 2023]. Moreover, popular Abesuriya et al.’s model (2017) for EEG and Deco at al’s (2017) Hopf oscillator based model were referred for comparison (see Fig. 9.(b) in main text).

##### S1.5.1 The Network of Hopf Oscillators by Deco et al

In the model of Deco et al [Deco et al., 2017], the Hopf oscillators are connected by a diffusion type real-valued coupling with a global coupling factor(G), bifurcation parameters (*µ*) are optimized iteratively. However, we fixed the amplitude of the oscillators at near zero (near the bifurcation point) in this case. A schematic diagram is given in Fig. 9.(a) (on the left panel of the figure) in the main text. Network dynamics is described as follows-

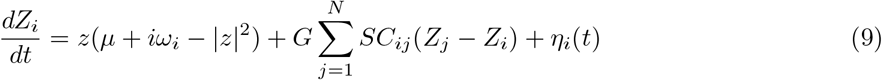

The complex form can be written also in Cartesian (*x, y*) or in polar coordinates(*r, θ*). The global coupling parameter, G is updated iteratively from 0.1 to 6, and fixed at an optimized value. The *µ* or bifurcation parameter is kept constant at around *µ* ≈ 0 and an additive Gaussian noise (AGN) is applied as *η*(*t*). For more details please refer to Deco et al [Deco et al., 2017].

For comparison purpose, 3 minutes 14 seconds of the Berlin dataset is used with number of oscillators, N= 68. The frequencies are fixed such a way that for a given oscillator (corresponding to a given ROI), the intrinsic frequency *ω_i_*, is set to be same as the frequency with the highest power obtained from the spectral domain data from that ROI. After simulation, the optimized G is found to be at 2.3, and the prediction power is at 0.34. Note that multiple parameter optimization procedures have been tried to optimize the parameters so that the best fit can be observed between simulated and empirical dataWischnewski et al. [2022].

##### S1.5.2 Neural Mass Model

The neural mass model is more biologically plausible, wherein a given neural mass consists of a balanced pair of excitatory and inhibitory neuron populations. Indeed, such a model is not a network of Hopf oscillators; however, they have been extensively used to study neural activity, reconstruct EEG data, and even the BOLD signal with approximating haemodynamic response function (HRF). The involved equations for constructing neural mass model are given below-

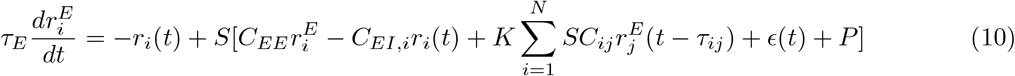

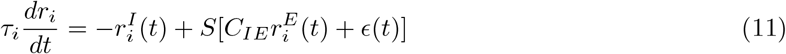

Here, equations 10 and 11 refer to the dynamics of excitatory and inhibitory neurons. Connections among the neural populations are shown in the Fig. 9.(a) in the main text with a schematic diagram [in the right panel of the figure]. The response function is given by S(x)-

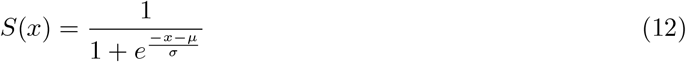

Readers are requested to consult the earlier works done by Deco et al. (2008), Abesuriya et al.(2018). to know more of those parameters [Deco et al., 2008, Abeysuriya et al., 2018]. Such a model was found to be extremely parameter sensitive, i.e constant input parameter, P, and the ratio between *τ_E_* and *τ_I_* often determine the system dynamics. Important to note that, *τ_E_* and *τ_I_* is scaled in such a way that the model gives the oscillation with 11*Hz* (approx) for Abesuriya et al.’s model and 40*Hz* for Castaldo et al’s model. Both Abesuriya et al.(2018) and Castaldo et al.(2023) used this model to simulate the neural activity that can generate EEG and BOLD signals individually. For simulating the Model 3, as given in Table 1 in the main text, the parameters given in the Castaldo et al’s. work was utilized to decipher the model’s accuracy to reconstruct EEG dataset from “Berlin” dataset. We have also used the model parameters given by Abesuriya et al. [Abeysuriya et al., 2018](Table 1 as given in that article) for simulating EEG dataset from “Berlin” dataset (see Fig. 9.(b) in the main text) to show the model’s performance for a different dataset to reconstruct EEG dataset.

Another important aspect of this model is the balance between the excitatory and inhibitory neurons’ populations. For that, the inhibitory connection is updated with a spike time-dependent process with a very small time constant to approximate the excitatory activity to a particular value *ρ* (Here, *ρ* and *τ_isp_* are fixed at 0.15 and 2.5). This process was referred to as homeostatic plasticity or inhibitory synaptic plasticity. The governing equation for homeostatic plasticity is given by –

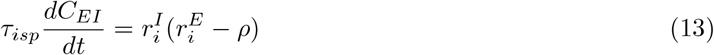

##### S1.5.3 The Network of Hopf Oscillators by Castaldo et al

In this study, for comparision, we also used the model of Castaldo et al.(2023) to simulate the EEG signals with Berlin dataset [Castaldo et al., 2023] (in this study). However, the model is almost similar to earlier Deco et al.’s model, but there are some striking differences. Firstly, the model includes time-delay, which is parametrized by the velocity value. Next, the *µ* is kept fixed at −5 eliciting damped oscillation.

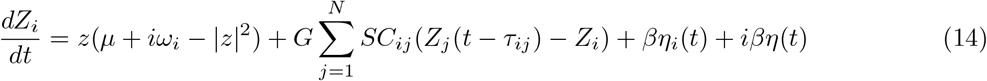

This model has two optimizable variables i.e *G* (global coupling), *< τ > orMD* (mean delay), which provide best approximation of FC, and FCD from empirical data. *η*(*t*) refers to the additive Gaussian noise (AGN). This model is used to reconstruct the empirical EEG data from Berlin dataset, of duration 3 minutes 14 seconds. The *G*, and *MD* parameters are updated iteratively. It was found that the model exhibits higher accuracy, and provides an intuitive understanding of incorporating the time delay parameter in whole brain model simulation.

### S2 Results

#### S2.1 Accuracy of the model

It has been demonstrated that the model simultaneously captures the BOLD and EEG signals with high accuracy. However, this accuracy, as we discussed, is strictly dependent on the model parameters, i.e. number of oscillators in each sub-cluster, number of learning epochs, number of hidden neurons in each layers. Interestingly, we have shown such importance of parameters on model’s capability in our earlier work [Bandyopadhyay et al., 2022].

However, in the main text, we discussed the effect of varying the number of oscillators in each sub-clusters, i.e. *n_HF O_* or *n_LF O_*. Tables S3 and S4 show the effect of varying *n_HF O_* or *n_LF O_* on the accuracy of empirical signal reconstruction, which can be expressed in terms of Pearson’s correlation coefficient (between simulated and empirical fMRI and EEG signals from a single ROI). Noteworthy, the result shown in the tables are for the first ROI’s simultaneous EEG-fMRI data from the third subject as indexed in “Berlin” dataset.

**Table S3:**
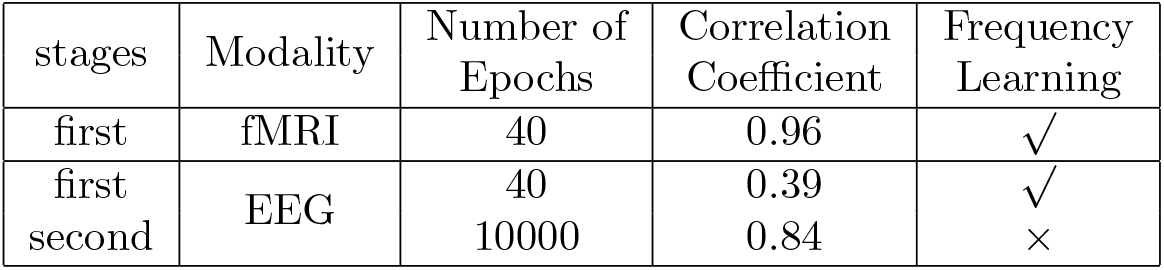
Performance of the model with *n_HF O_* = 5, *n_LF O_* = 5.

**Table S4:** Performance of the model with *n_HF O_* = 10, *n_HF O_* = 10.

| stages | Modality | Number of Epochs | Correlation Coefficient | Frequency Learning |
| --- | --- | --- | --- | --- |
| first | fMRI | 40 | 0.98 | ✓ |
| first | EEG | 40 | 0.52 | ✓ |
| second |  | 10000 | 0.97 | × |

#### S2.2 Structural Disconnection Based on Absolute Thresholding

In the main text, the impact of proportional threshold was described. However, we have not discussed the impact of absolute threshold. For absolute threshold, a certain value of structural connectivity 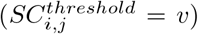 was picked and the SC was pruned such a way that, *SC_i,j_* = 0, where*, SC_i,j_ <*= *v*. Interesting to note that, with increasing *v*, the model loses its capacity to capture the empirical functional connectivity map (FC, FCD) along with the modular architecture of brain network for both EEG and fMRI data.

##### S2.2.1 Impact on FC

Impact of structural connectivity on Functional connectivity, can be estimated by evaluating the simulated FC from SC, which is pruned by absolute threshold. In this case, we fixed the threshold value, *v* at 0.05, 0.10, and 0.20. It is found that the correlation value or predictive power between simulated FC and empirical FC keeps decreasing with increasing *v*. For electro-physiological data this is shown in the Fig. S1a. For electrophysiological data, for all the band-patterns, the FC prediction deteriorates with increasing values of threshold parameter, *v*. A similar outcome can be observed in case of BOLD signals, which was shown in Fig. S2a.

##### S2.2.2 Impact on FCD

FCD is typically estimated by taking a sliding window technique, where each segment of signals (by taking a window with fixed length) from all the ROI is transformed into FC matrix (forming a *K* ×*K* ×*N_t_* tensor, where *K* is number of ROIs and *N_t_* is the number of windows) by either taking Pearson’s correlation or Amplitude envelope correlation (AEC). For simplification, the FCD matrix is built by taking Pearson’s correlation between *N_t_* number of FC matrices, yielding a *N_t_*×*N_t_* matrix. For understanding the impact of pruning, FCD is simulated from the model with prunned SC at different level as described earlier. Later, empirical and simulated FCD is compared with Kolmogorov-Smirnov distance (*KS_distance_*) and given in the Figs.S1b and S2b. A Similar analysis was done to understand the impact of structural pruning by proportional threshold on FCD also (shown in main text). The Fig. S1b illustrates the impact of FCD by increasing the threshold value, *v*, which elicits that the distance between the distribution of FCD values between simulated and empirical FCD. *KS_distance_* increases monotonically with increasing value, *v*. Figure S2b reveals that the FCD arising from the BOLD signal pursues the similar trend, referring elevation of *KS_distance_* with increase in *v* value.

##### S2.2.3 The Impact on Community Structure

Louvain algorithm was used for community detection; however, this algorithm has the problem of degeneracy. A symmetric matrix *D_i,j_* is evaluated representing the community structure, where each element *i, j* represents the probability of an ROI/node affiliated to a particular community resulting from multiple iterations of the algorithm as stated in the main text. We showed that such a study can decipher the impact of structural loss in the aberration of modular architecture in network emerged from functional Both in experimental and theoretical studies, a loss of modularity in the network architecture, as obtained from EEG data, has been observed earlier. We showed that such a loss is reflected in both modular architectures in brain network computed from EEG (for all the prominent frequency bands-*α, β, δ, θ, γ*) and fMRI data within a single mathematical framework (see the Fig. S2c). Note that, in the main text, only the modular community structure of *α* band EEG is computed and used for comparison between model’s output from pruned and before-pruned SC. Figure S1c shows that with increasing *v*, the community structure deviates from its original (before pruning) condition. This is measured by computing the correlation value between the simulated and empirical community structure matrix, and the difference in correlation value for different pruning conditions marks the deviation in the community structure.

**Figure S1:**
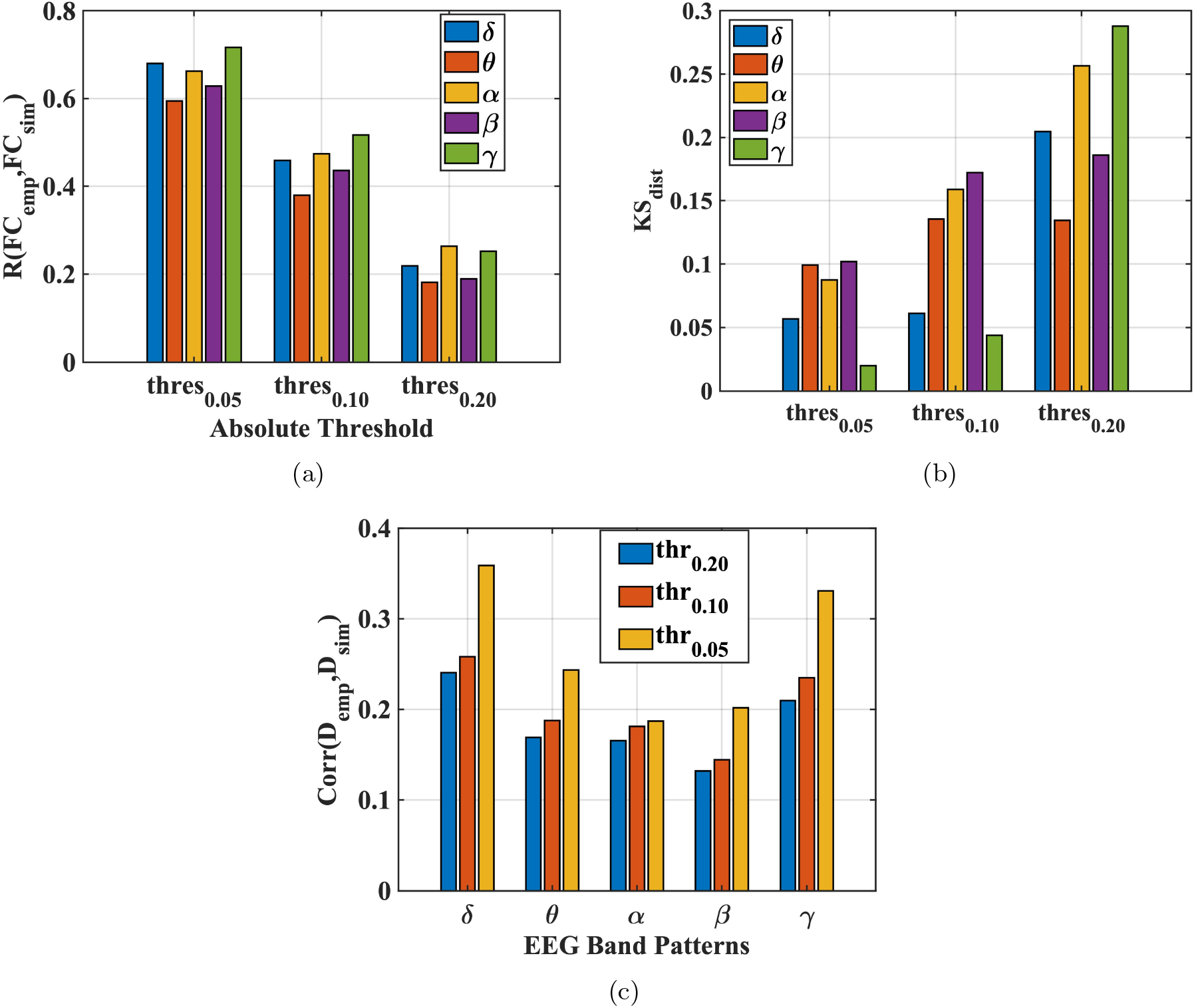
Impact on the functional connectivity map due to structural pruning for electro-physiological data. Fig.(a) represents the loss of FC due to structural pruning for all the frequency bands. Figure (b) shows the impact of pruning on FCD in terms of *KS_distance_*, and Fig. (c) reveals the community structure simulated from the model after perturbation with different absolute threshold values, *v*. data. Similar to the case of proportional threshold, with the increasing absolute threshold, *v*, the model can not approximate the community structure.

##### S2.2.4 Spatio-Temporal Integration

The brain dynamics are characterized by a delicate balance between integration and segregation, a balance that can be disturbed by certain perturbations. In the proposed computational study, we showed the impact of structural loss on integration of functional brain network, and showed that it reduced in case of perturbation by thresholding structural connectivity - both absolute and proportional thresholding.

Figures S3a, and S3b illustrate that the model can not retain the integration level after structural loss. Such an in-silico perturbation study will help to understand the loss of integration in neurological and cognitive disorders. Figure S3a shows the loss of integration in terms of ISOR (Integration state Occurrence Rate), where the predominantly integrated states are calculated as described in detail in the main text. This is done for analyzing ultra-slow haemodynamic signal. Interestingly, for the electro-physiological signal, the integration level (*M_in_*) is calculated with Nested Spectral clustering (NSP), from the spectral graph theory [Zhou et al., 2023]. The results show that for all the frequency bands, the integration level reduces with increasing threshold parameter, *v*.

**Figure S2:**
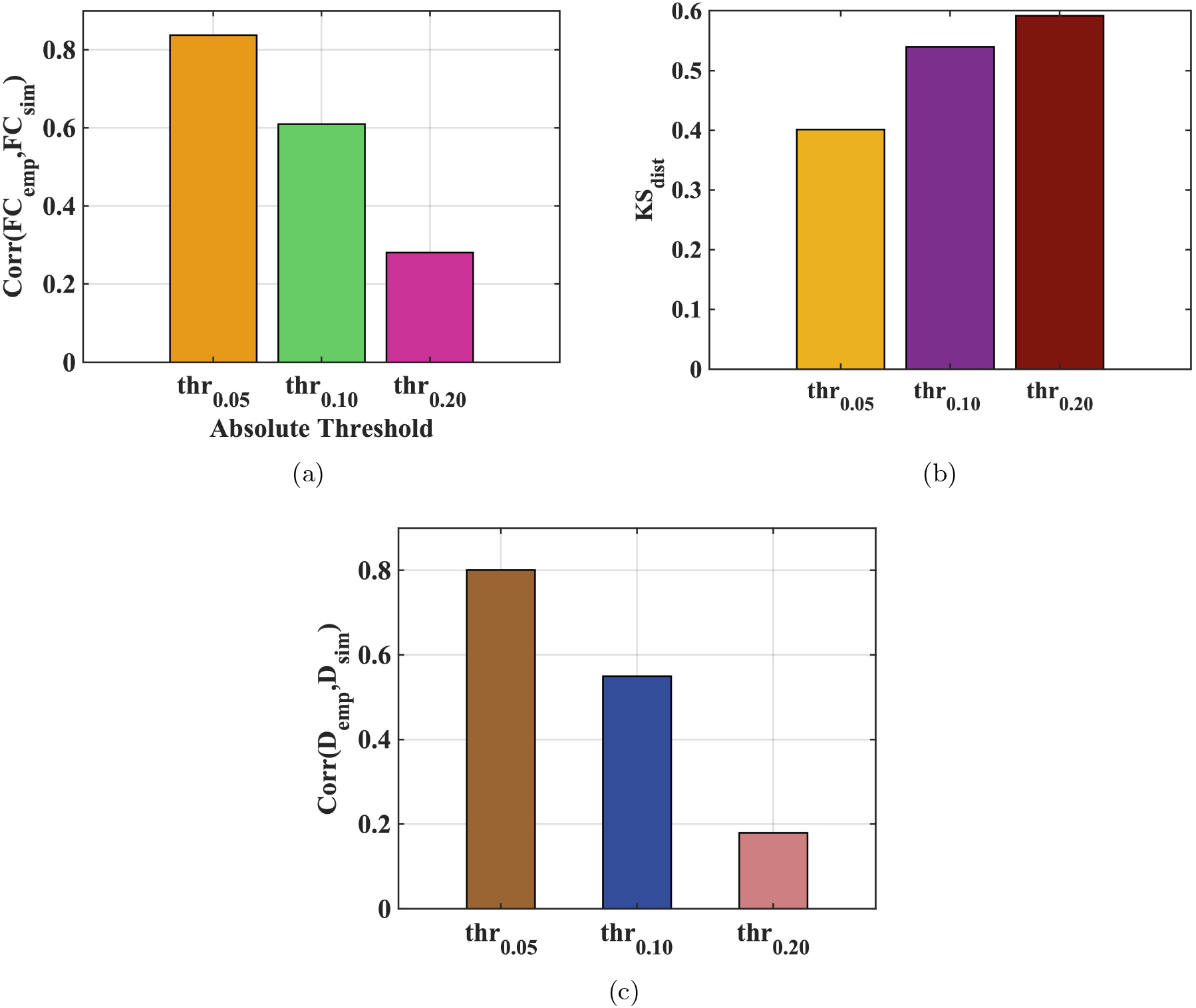
The Impact on functional connectivity map along with the modular architecture due to structural pruning in case of BOLD signals. Figure (a) represents the loss of FC due to structural pruning for all the frequency bands, and Fig.(b) shows the impact of pruning in terms of *KS_distance_*. Figure (c) reveals the community structure simulated from the model for different absolute threshold values, *v*.

From this in-silico perturbation study, we conclude that the impact of FC, FCD, community structure, spatio-temporal integration pattern by structural pruning is not constrained by the approach of pruning procedure and the aberration of SC can cause multifaceted impact on spatio-temporal brain dynamics.

**Figure S3:**
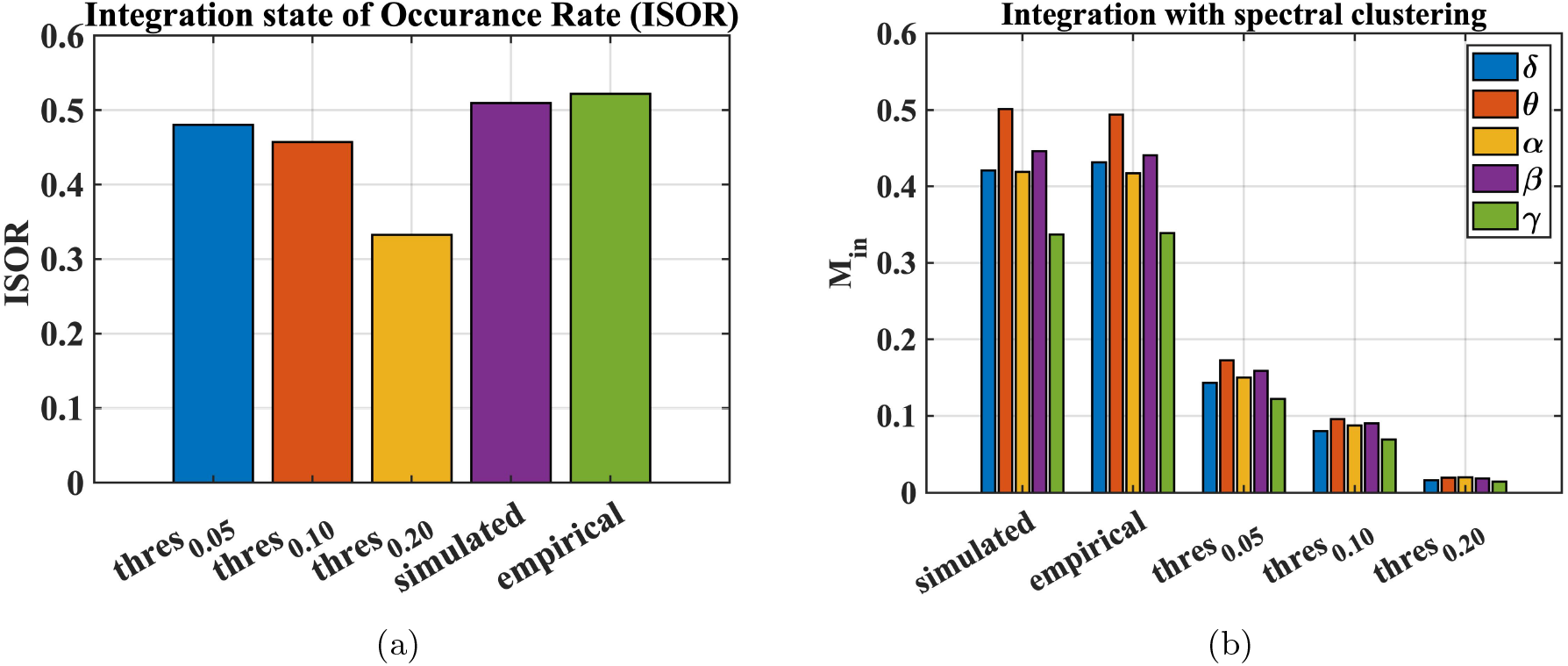
Evaluation of integration from functional connectivity network for different structural pruning. Figure (a) shows the impact of pruning on integration level of brain in case of haemodynamic signal, and for electrophysiological signal. The methodological details involved here can be found in the main text.

## Notes

### Competing Interest Statement

The authors have declared no competing interest.

